# Post-acquisition super resolution for cryo-electron microscopy

**DOI:** 10.1101/2023.06.09.544325

**Authors:** Raymond N. Burton-Smith, Kazuyoshi Murata

## Abstract

Super resolution detector acquisition for cryo-EM has been used to improve the clarity of cryo-EM reconstructions. Recent reports have demonstrated achieving resolutions beyond the physical Nyquist limit using super resolution acquisition. Here, we demonstrate exceeding the physical Nyquist limitation by pre-processing the raw micrograph movies from “counting mode” data which has already reached physical Nyquist reconstruction resolution. To demonstrate functionality, micrograph movies of five datasets were pre-processed and demonstrate that it is possible to exceed the physical Nyquist limit via pixel doubling before motion correction. We call this “post-acquisition super resolution”, or PASR. While this was originally developed for processing of giant virus datasets, where acquiring at high magnification is not always possible or desirable, it is also shown to work for smaller objects such as adeno-associated virus (AAV) and apoferritin, both of which are still high symmetry, and jack bean urease, with lower symmetry. PASR could reduce the magnification required to achieve desired resolutions, which may increase collection efficiency. PASR can also be of use for in vivo tomography and facilities with high storage demands. However, this method should only be used for data which is able to achieve the Nyquist limit without PASR pre-processing. It will not improve attained resolutions of data which does not already reach the Nyquist limit.

## Introduction

In cryo-electron microscopy (cryo-EM), while higher magnification does not always mean higher resolution(Kayama et al., 2021), the magnification at which data were acquired always causes a hard (unbreakable) limit on the maximum resolution attainable with a dataset. This is derived from the work of H. Nyquist (Nyquist, 1928) and C. Shannon (Shannon, 1949) at Bell Laboratories for electronic communications, and is applicable to any digital signal processing. These theories, often called the Nyquist theorem or Shannon theorem (or a combination of the two) is, in cryo-EM, frequently called the “Nyquist limit”. In brief, the Nyquist limit is half the sampling rate of the detector (or the reciprocal of double the pixel size of a digital image) and is the absolute theoretical limit on resolution. Normally, however, other factors come into play before the Nyquist limit to limit the resolution of cryo-EM images, such as the microscope environment and sample condition. Nonetheless, if this limit is reached, increasing the resolution of cryo-EM 3D reconstructions traditionally required the collection of a dataset at higher magnification. We recently demonstrated reaching this limit with Melbournevirus (Burton-Smith et al., 2021) and the same effect (reaching the Nyquist limit) will have been observed by any cryo-EM user who has downsampled/binned their data to speed earlier stages of processing.

Direct electron detectors (DEDs) are near ubiquitous in cryo-EM facilities now, having gained in popularity due to their high-resolution capabilities and enhancing acquisition automation. DEDs contributed to the “resolution revolution” (Kühlbrandt, 2014) along with GPU acceleration and other software developments and more stable microscopes. This created an explosion of interest in cryo-EM (Iudin et al., 2023). Early DEDs functioned similarly to scintillator coupled CCDs which had previously been popular, with a “linear” or integrating recording mode (Fig. 1a). With the release of the Gatan K2 Summit (AMETEK, USA), “counting mode” became available, where the controller estimates where the electron originally interacted with the sensor and defines that pixel as the whole signal (Fig. 1b) and finally, super resolution acquisition (Fig. 1c) where the electron is localized to a quadrant of a pixel. Along with dose fractionation, allowing the acquisition of “movies” of cryo-EM data to compensate for stage instability and particle movement (Brilot et al., 2012; Campbell et al., 2015), direct detectors catapulted achievable resolution forward.

**Figure 1:**
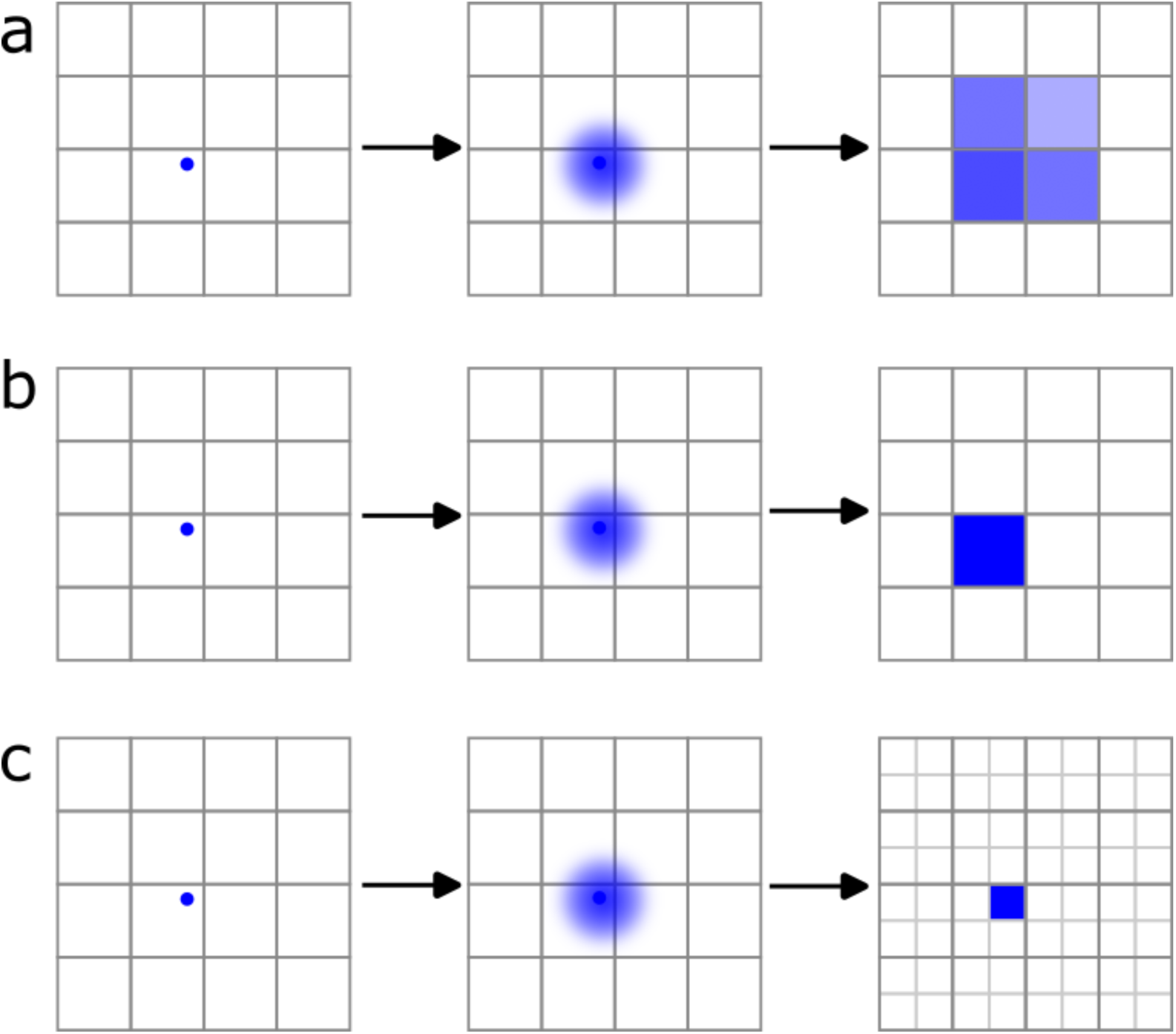
Diagrammatic representation of electron detection on a modern cryo-EM direct detector. (a) Integrating (or linear) mode; in this mode, the detector acts in a similar way to old scintillator-coupled CCD devices. When an electron strikes a pixel (left), a so-called “electron event”, the charge diffuses through the silicon (centre) such that multiple pixels are read out by the controller with a partial signal from each. This makes the image more blurred, (b) counting mode; in this mode, upon an electron event the controller calculates the pixel with the greatest signal contribution and assigns all the “event” to that single pixel, (c) super resolution mode; this is an enhanced version of counting mode, where the controller estimates where the electron first interacted with a pixel with sub-pixel (quadrant) accuracy.

Super resolution is a technique now widely used across a range of disciplines for acquisition of data, although implementations vary (Betzig & Trautman, 1992; Chiu et al., 2015; Guerra, 1995; Poot et al., 2013). In cryo-EM, the first commercially available camera/detector with super resolution capability was the Gatan K2 Summit, which is now available in direct detectors from Gatan (Pleasanton, Ca., USA), Thermo Fisher Scientific (Waltham, Ma., USA) and Direct Electron (San Diego, Ca., USA), but it is not ubiquitous. The technique involves algorithmic estimation of an electron event (where an electron strikes the detector in such a way as to be recorded) in a pixel sub-section (Fig. 1c), but it must be explicitly selected at acquisition time. A great deal of use has been made of super resolution acquisition since it became available to cryo-EM facilities, and many super resolution datasets are available publicly in the EMPIAR database (Iudin et al., 2016; Iudin et al., 2023). Recently reconstructions “past Nyquist” have been demonstrated using super resolution data (Feathers et al., 2021) – although this data was still explicitly recorded at acquisition time with super resolution mode. There has also been recent interest in neural network optimisation similar in effect to our work presented here (Huang et al., 2023).

Giant structures (particles >150 nm) present a significant challenge to cryo-electron microscopy. Resolution limitations caused by defocus gradients can be compensated for by using Ewald sphere correction (DeRosier, 2000; Russo & Henderson, 2018; Wolf et al., 2006; Zivanov et al., 2018) or “block-based” reconstruction methods (Zhu et al., 2018) or a combination of the two. Recently, Ewald sphere correction has been demonstrated to improve resolution of objects as small as apoferritin (Nakane et al., 2020; Yip et al., 2020). Ultimately, however, for giant structures the object of interest is extremely large for high resolution single particle reconstruction (SPR) cryo-EM due to difficulties in data acquisition.

As a result, data acquisition for giant structures is a careful balancing act between a high enough magnification to achieve (near) atomic resolution and low enough magnification to avoid needing to collect tens or hundreds of thousands of micrographs to acquire a similar number of usable particles. One previous work on giant viruses demonstrated the need to acquire almost one micrograph per selected particle (Wang et al., 2019).

We recently reported (Burton-Smith et al., 2021) a reconstruction of Melbournevirus, a giant virus with a maximum dimension of ~250 nm, to a resolution of 4.9 Å for the whole viral particle and to 4.42 Å (the maximum attainable resolution from that dataset) for block-based reconstructions of the capsid at the two-, three- and five-fold axes. Subsequently, however, we were able to apply Bayesian polishing (Zivanov et al., 2019) to the whole virus to reach the Nyquist limit for the whole virus reconstruction (unpublished data). Conversely, we have also reported the use of super resolution acquisition for Tokyovirus, but which did not reach the physical Nyquist limit, and thus super resolution may have been detrimental instead, due to the lower detective quantum efficiency (DQE) we observed at 1 MV (Chihara et al., 2022).

As reported, when using the block-based reconstruction method for Melbournevirus, after localised defocus refinement the solvent-corrected gold-standard FSC curves reached the Nyquist limit above the 0.143 metric, indicating that if data had been able to be collected at higher magnification, higher resolution would have been achieved. This caused us to ponder: would pre-processing of the counting mode data to simulate “super resolution” data permit us to break this resolution limit as super resolution at acquisition time has done? Would it cause artefacting in the resulting reconstruction? In short; yes, it does, and no, it does not, respectively.

Here, we demonstrate the potential of a “post-acquisition” super resolution-like pre-processing step, where each pixel of each frame of a counting mode micrograph (Fig. 2a) is split into four identical sub-pixels before motion correction (Fig. 2b). After motion correction is performed, each pixel in the resulting sum is unique (Fig. 2c). This is the “dithering” method, first used in astrophotography (Adorf & Hook, 1995; Hook & Fruchter, 2000) applied to cryo-EM. As such, we require at least a small amount of motion between frames, but of course too much movement becomes detrimental. This does not require complex multi-pass neural network processing to achieve resolution improvements like cryo-ZSSR (Huang et al., 2023).

**Figure 2:**
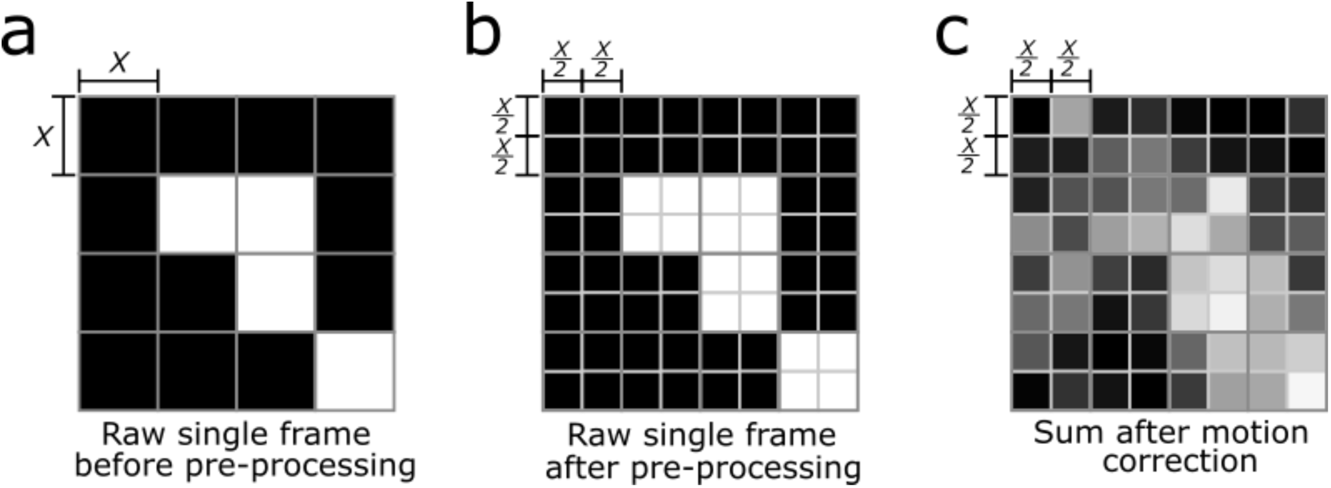
Diagrammatic representation of the post-acquisition super resolution (PASR) method. (a) a normal micrograph frame where each pixel has the effective magnification of *x*×*x* Å. This is pre-processed using a script in GNU Octave to make each pixel a 2×2 grid of ½*x*×½*x* Å (b) across all pixels and all frames of the micrograph. Upon motion correction, each sub-pixel in the resulting micrograph is unique (c).

As this effectively simulates super resolution without the sub-pixel localisation, but rather dithering during frame alignment, we call this “post-acquisition super resolution”, or PASR. We demonstrate this for two publicly available datasets of higher and lower symmetry: apoferritin, an octahedral symmetry complex which is now a favourite among the cryo-EM field for benchmarking and testing (Danev et al., 2019; Hamaguchi et al., 2019; Kayama et al., 2021; Nakane et al., 2020; Noble et al., 2018) and which currently holds the world record for attained cryo-EM resolution (Yip et al., 2020), and jack bean urease (Feathers et al., 2021) as it is lower symmetry, and a public dataset was available on EMPIAR (Iudin et al., 2016; Iudin et al., 2023) which allowed us to compare between the results of super resolution acquisition and our PASR technique, and also permitted testing for 200 kV data. Three other datasets of higher symmetry were tested also; first, an adeno-associated virus (AAV) dataset generously provided by a collaborator from a different study, which allowed us to test the application of PASR to correlative double sampling (CDS) acquisition. Second, a subset of an apoferritin dataset acquired at nearly 2 Å/pixel, and third, by reprocessing our previously reported Melbournevirus dataset (Burton-Smith et al., 2021) which inspired this work. We further carry out a brief demonstration of the PASR method on already motion corrected micrographs, albeit with inferior results compared to application to “movies” prior to motion correction, which is as we expected.

## Results

### Motion correction and CTF estimation

We tested motion correction with both MotionCor2 (Zheng et al., 2017) and the RELION implementation (Zivanov et al., 2018), both of which proceeded normally, with no errors. A high-zoom visual examination of motion corrected micrographs using Fiji/ImageJ2 (Schindelin et al., 2012) revealed no abnormalities, with individual pixels differing from neighbouring pixels. We tested datasets from a range of detectors and accelerating voltages to see whether acquisition hardware affected the effectiveness of PASR (Table 1).

**Table 1:** Datasets used in this work.

| Dataset | Apoferitin (Apo64K) | Apoferitin | Melbournevirus | Adeno-associated virus | Jackbean urease | Nodavirus |
| --- | --- | --- | --- | --- | --- | --- |
| Source | This study | EMPIAR-10216 | Burton-Smith *et al.* (2021) | This study | EMPIAR-10549 | EMPIAR-10203 |
| Physical pixel size (Å/pixel) | 1.91 Å/pixel | 0.517 Å/pixel | 2.21 Å/pixel | 1.34 Å/pixel | 2.1 Å/pixel (super res: 1.05 Å/pixel) | 1.06 Å/pixel |
| Microscope | Titan Krios G4 (300 kV) | Titan Krios G3 (300 kV) | Titan Krios G2 (300 kV) | CRYOARM 300 (300 kV) | Tecnai Arctica (200 kV) | Titan Krios (300 kV) |
| Detector | Falcon 4 | Falcon 3 | K2 Summit | K3 | K3 | K2 Summit |
| Acquisition Mode | Counting (EER) | Counting | Counting | CDS | Super Resolution | Counting |
| Format | EER, converted to TIFF (4K/8K) | MRC (gain corrected) | MRC (gain corrected) | TIFF (movie, separate gain) | TIFF (movie, separate gain) | MRC (single frame) |
| Movies or micrographs | Movies | Movies | Movies | Movies | Movies | Micrographs |
| Processing carried out | 4K, 8K, PASR | Original, binned, PASR | PASR | Original, PASR | Original, binned, PASR | Original, PASR |
| Relevant Figures | 7, S1-5 | 5, S14-16 | 8, S6-8 | S9-10 | 6, S11-13 | S18-20 |

If we were to encounter issues, CTF estimation was expected to be the first point of the processing pipeline to exhibit it. Figure 3 shows power spectra from the same original apoferritin micrograph from dataset 1 (hereafter, “Apo64K”) when processed at 4K (native sampling) (Fig. S1), 8K super resolution (Fig. S2) and 8K PASR (Fig. S3) settings (Table 2). Underneath each power spectrum in Figure 3 is the 1D diagnostic plot from CTFFIND 4.1.14 (Rohou & Grigorieff, 2015), showing that estimated defocus values and azimuth are comparable between the three. The 4K micrograph power spectrum appears zoomed as the sampling frequency limit is 3.82 Å rather than 1.91 Å as is the case for super resolution and PASR samplings. Occasionally, a few micrographs “fail” CTF estimation – where the simulated CTF fits the Thon rings poorly – with PASR processed but not in the original data (see Tables 2–7). This may be caused by the decrease in contrast (Fig. S1-3a) for super resolution and PASR micrographs relative to micrographs acquired at physical Nyquist sampling. Micrographs which “pass” CTF estimation demonstrate very similar defocus, astigmatism, and maximum resolution estimates (Fig. 3, Table 2–7). Only CTFFIND4 (Rohou & Grigorieff, 2015) was used for RELION. On the same micrograph, the patch CTF estimation of CryoSPARC (Punjani et al., 2017) reports similar values to CTFFIND for the 8K super resolution and 8K PASR data (Figs. S4C and S5C).

**Figure 3:**
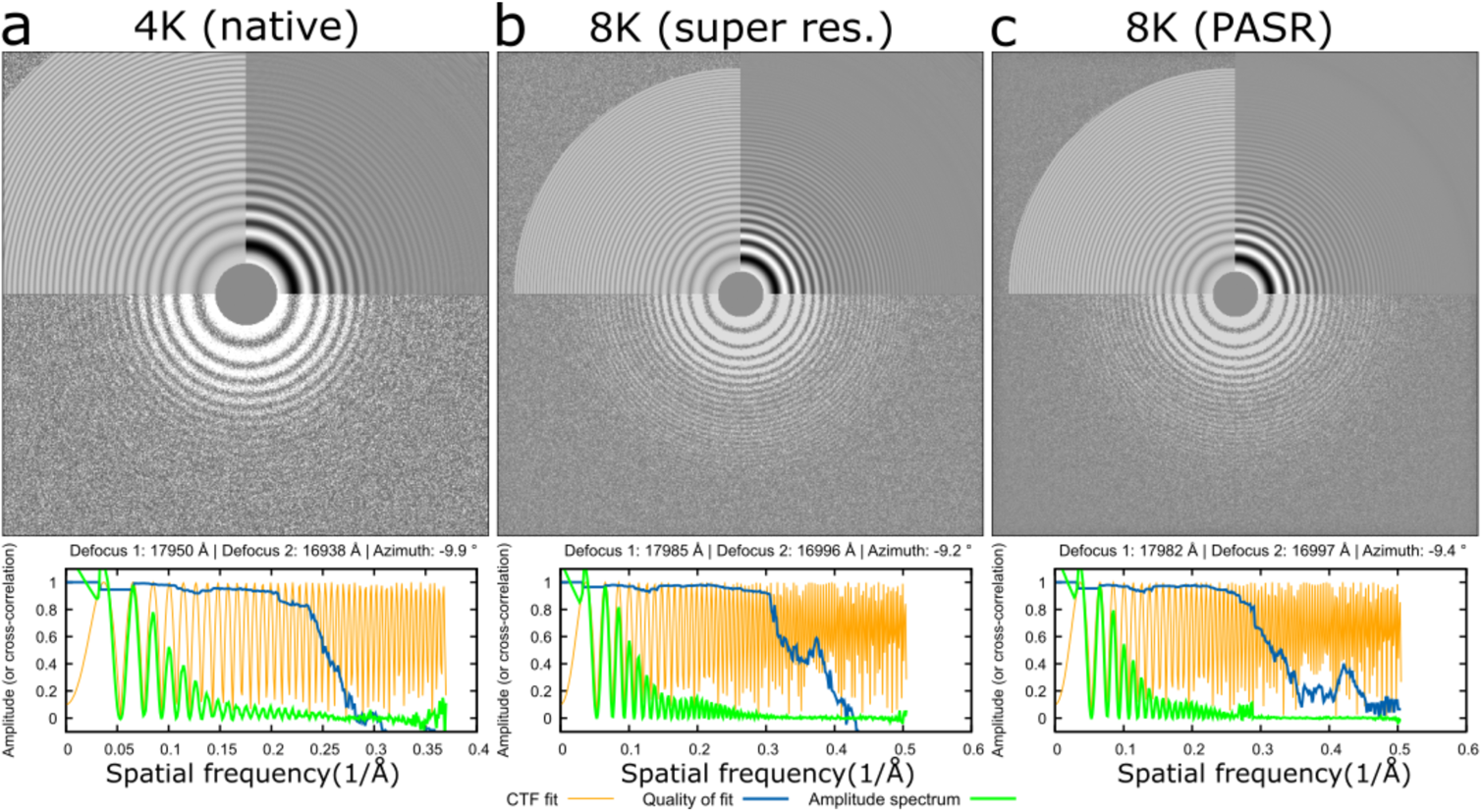
Comparison of contrast transfer function (CTF) estimation for 4K (native), 8K (super resolution) and 8K (PASR-preprocessed 4K micrographs) from the same micrograph. The original micrograph was collected using the EER file format and converted to either 4K TIFF or 8K TIFF using the RELION function relion_convert_to_tiff. CTFFIND was used to calculate parameters for each with the following settings: FFT box size: 512, Minimum resolution: 30 Å, Maximum resolution: 3 Å, Minimum defocus: 1000 Å, Maximum defocus: 50,000 Å, Defocus step size: 100 Å. For each condition, the power spectrum is displayed, showing clear Thon rings, along with the CTFFIND derived rotational average and simulated fit (top). Underneath, 1D diagnostic plots from CTFFIND are shown, demonstrating highly similar calculated defocus and astigmatism values. The sampling frequency of the 4K micrograph is 1.912 Å/pixel, while for the super resolution and PASR micrographs sampling frequency is 0.956 Å/pixel.

**Table 2:** Parameters for apoferritin processing for small dataset at low magnification using RELION.

| Apoferritin (this study) (RELION 4) |  |  |  |
| --- | --- | --- | --- |
|  | Original (4K) | Original (8K) | PASR |
| Accelerating voltage | 300 | 300 | 300 |
| Spherical aberration | 2.7 | 2.7 | 2.7 |
| Micrographs (total) | 32 | 32 | 32 |
| Pixelsize (Å/pixel) | 1.912 | 0.956 | 0.956 |
| Passed CTF estimation | 32 | 32 | 32 |
| CTF estimation cutoff (Å) | n/a | n/a | n/a |
| Particles picked | 102,364 | 84,586 | 102,493 |
| Particles selected | 33,468 | 63,313 | 64,597 |
| Final resolution (Å) | 3.824 | 2.13 | 2.33 |
| Final particle count | 33,468 | 63,313 | 64,597 |
| Estimated b-factor | -116.6 | -62.4 | -99 |

### Particle picking and 2D classification

Laplacian-of-Gaussian (Zivanov et al., 2018) and template-based (Scheres, 2015) autopicking methods were unaffected (Figs. S6-13) by PASR processing, resulting in similar numbers of particles when used for a dataset (Tables 2–7) with the same parameters.

2D classification appears unaffected by PASR processing. Fig. 4 shows some example classes from different datasets and PASR is visually indistinguishable from native 2D classes. This is as expected as early stages of classification are usually carried out on downsampled data. Interestingly, the PASR data for apoferritin (EMPIAR-10216) (Table 3) classified into far fewer classes (Fig. S14) than the binned or original data (Fig. S15, S16) when using the same settings, but the number of particles contained within these good classes was approximately the same (Table 3) although we have not yet tested whether this is a repeatable phenomenon. Conversely, for the Apo64K dataset (Table 2, Fig. S1), the 4K sampling data had fewer clear classes and contained fewer particles in total than either the 8K sampling (Fig. S2) or PASR (Fig. S3) datasets.

**Figure 4:**
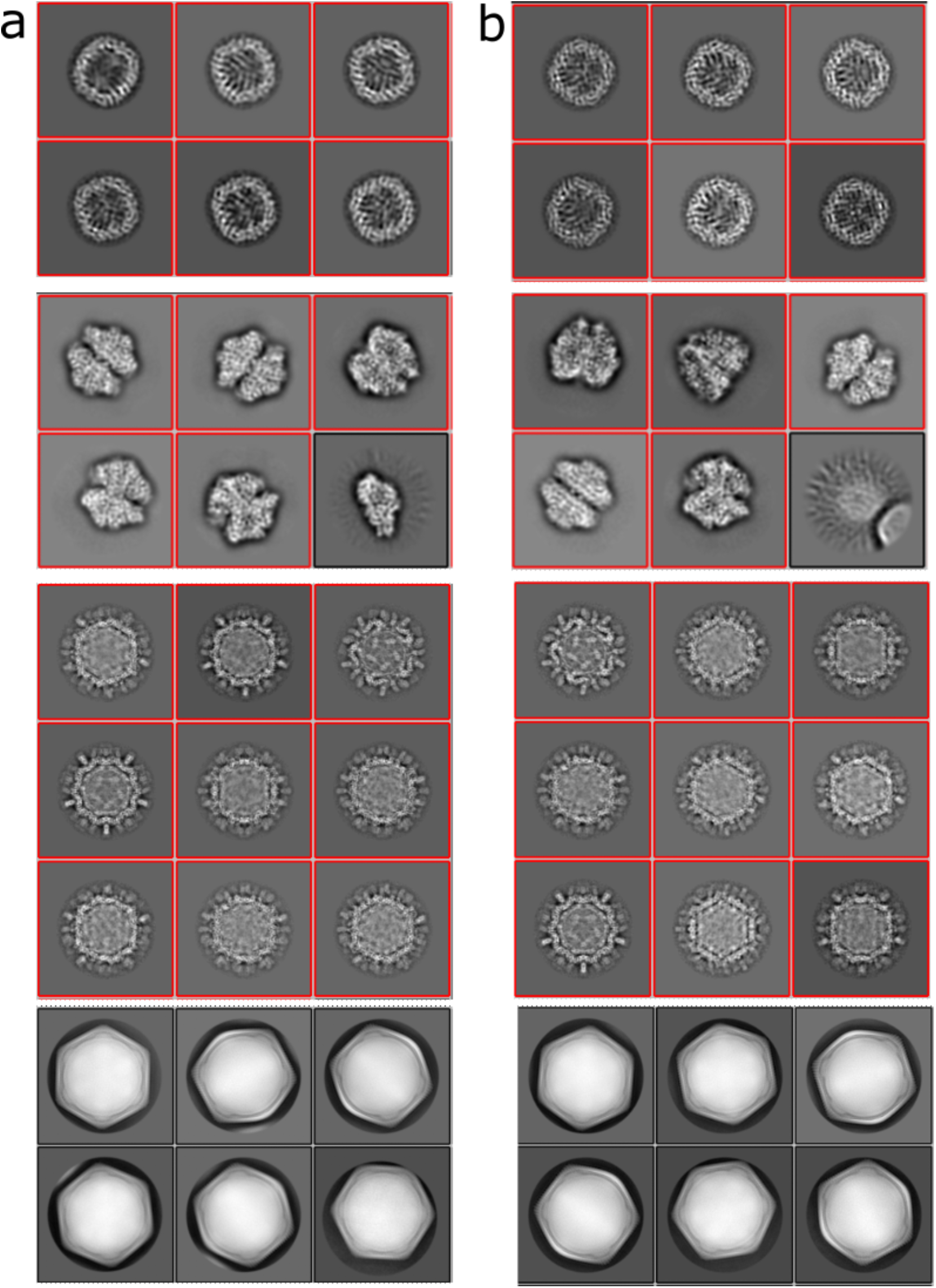
Example 2D classes from different datasets showing (a) classes from original data and (b) classes from PASR data. Classes are from apoferritin, jack bean urease, Nodavirus and Melbournevirus processing. Particles from different datasets not to the same scale.

**Table 3:** Parameters for apoferritin (EMPIAR-10216) processing.

| Apoferritin (EMPIAR-10216) (RELION 3.1) |  |  |  |
| --- | --- | --- | --- |
|  | Original | Binned | PASR |
| Accelerating voltage | 300 | 300 | 300 |
| Spherical aberration | 2.7 | 2.7 | 2.7 |
| Micrographs (total) | 1,647 | 1,647 | 1,647 |
| Magnification (Å/pixel) | 0.517 | 1.034 | 0.517 |
| Passed CTF estimation | 1,228 | 1,153 | 1,106 |
| CTF estimation cutoff (Å) | 5 | 5 | 5 |
| Particles picked | 217,495 | 156,099 | 207,388 |
| Particles selected | 161,966 | 149,536 | 152,367 |
| Final resolution (Å) | 1.71 | 2.068 | 1.78 |
| Final particle count | 161,966 | 149,536 | 152,367 |
| Estimated b-factor | -44.4 | -46.8 | -50.2 |

### Initial model generation

Initial model generation remains one of the more challenging aspects of cryo-EM single particle analysis (Gomez-Blanco et al., 2019; Grant et al., 2018; Kimanius et al., 2021; Punjani et al., 2017; Reboul et al., 2018; Zivanov et al., 2018). Much effort has been put into improving initial model generation but there is still some element of luck. We tested initial model generation throughout the datasets. For every case except one, the RELION initial model algorithm was successful in generating an acceptable initial model which resembled the 2D classes used. For the urease dataset (Table 4), the PASR processed dataset failed to generate a good initial model in RELION. To generate a good initial model from the PASR dataset we exported the particle stack to cisTEM (Grant et al., 2018) to successfully generate an initial model. As we have previously had success with cisTEM initial model generation where RELION initial model generation fails for “normal” data, while this must be noted we do not consider it an impediment to PASR. Initial model generation with the Variable-metric Gradient Descent with Adaptive Moments (VDAM) algorithm of RELION 4 (Kimanius et al., 2021) was successful with the 8K PASR apoferritin dataset but needed to be carried out a second time on the 8K super resolution dataset before a good initial model was generated. Initial model generation in CryoSPARC was successful on the first attempt for the PASR data but the second attempt for the super resolution data with default parameters, whether symmetry is imposed or not during initial model generation. With parameters better optimised for apoferritin (20 Å starting resolution, 8 Å final resolution, in the same manner as in cisTEM) initial model generation was successful first time for both.

**Table 4:**
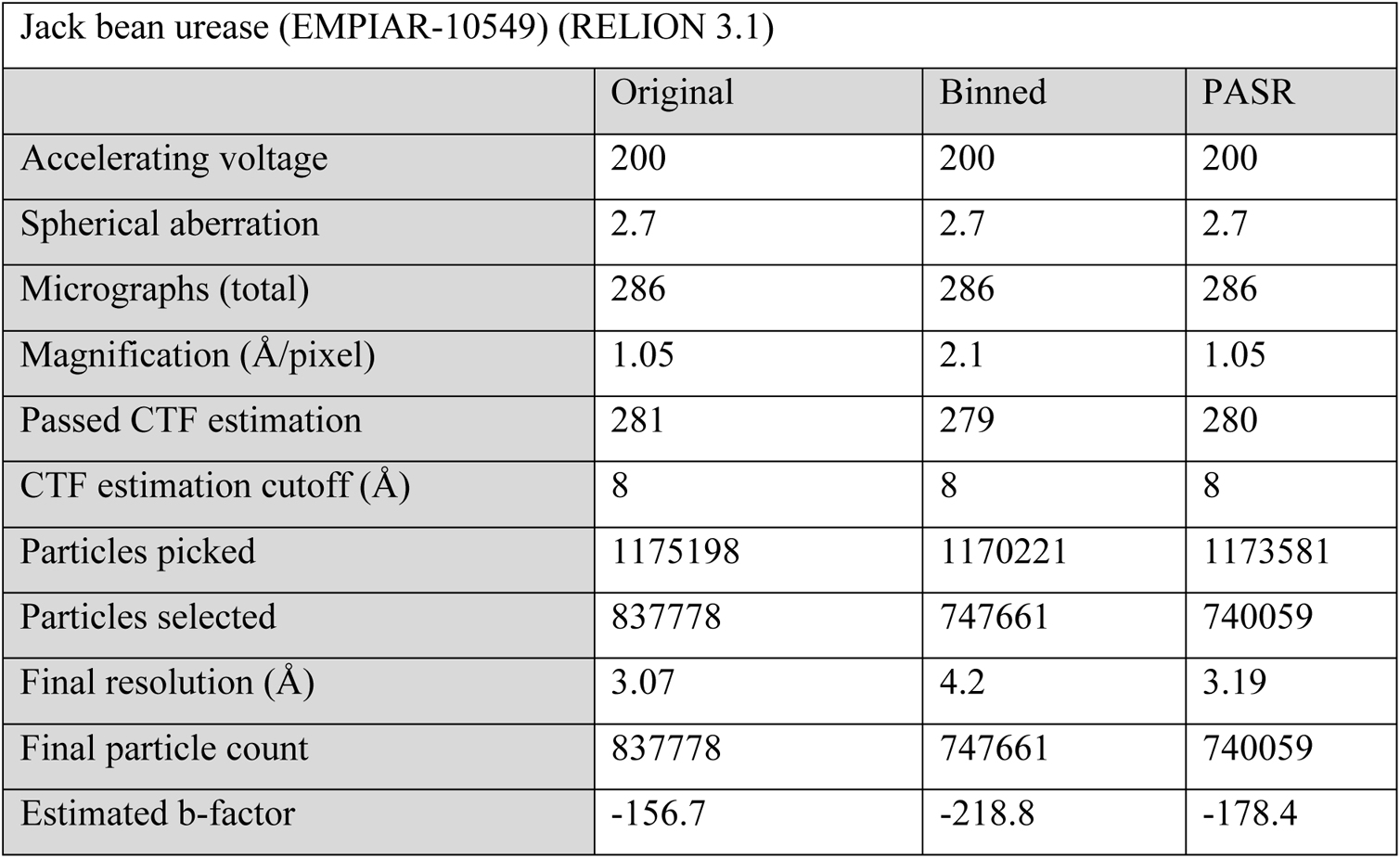
Parameters for jack bean urease (EMPIAR-10549) processing.

### 3D classification and imaging parameter refinements

3D classification (when used, e.g., Melbournevirus) and refinement displayed no anomalies in processing. Defocus and astigmatism refinement were unaffected, with refined values within expected tolerances given those obtained with the original datasets. Magnification anisotropy and beam tilt estimation were also within tolerance. Bayesian polishing likewise showed no adverse effects, although like native super resolution data, the PASR datasets took longer to analyse in the first step due to the increased size of the micrographs.

### Final map evaluation

PASR data suffers a marginal loss in final resolution compared to the original data for apoferritin (EMPIAR-10216) (Table 3) and urease (EMPIAR-10549) (Table 4), although this may be in part because of the need to first downsample the data from the original data uploaded to EMPIAR. However, the side chains look comparable (Fig. 5, 6). While PASR alignments report slightly worse resolution than the 8K sampled micrographs for our apoferritin dataset, once again, the side chains look comparable (Fig. 7). For both Melbournevirus (Fig. 8) (Table 5) and AAV (Table 6), PASR data is a visible improvement over those original data. Local resolution estimation of the PASR Melbournevirus blocks exceeds the Nyquist limit of the original data and is in line with local resolution estimations to be expected given the gold standard FSC reported (Fig. 8a). Other PASR processed datasets show the same when compared to Nyquist limited data (Figs. S1-16).

**Figure 5:**
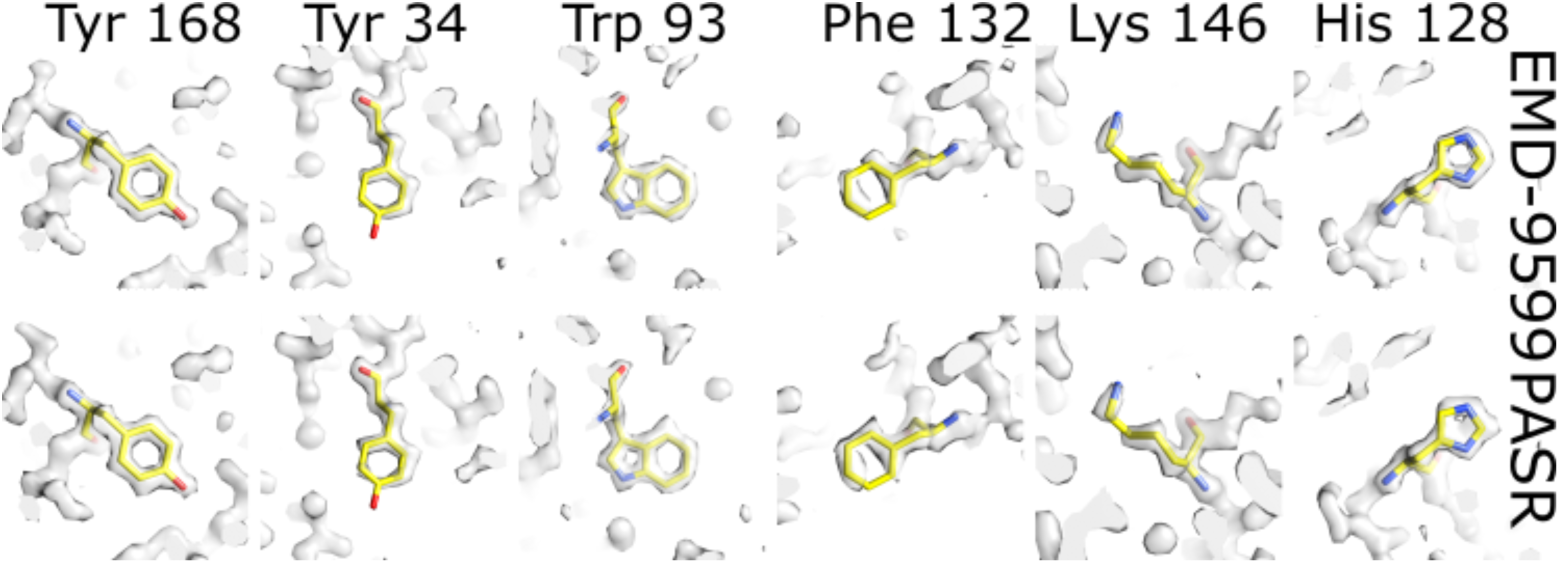
comparing side chains of PASR processed apoferritin data achieving <2 Å and directly comparing with EMD-9599 at 1.63 Å. When reprocessing EMPIAR 10216, 1.71 Å was achieved while PASR achieved 1.78 Å. Six example side chains of varying quality are shown; In EMD-9599, the five-member-ring nature of His128 is clear, while with PASR the hole is much less well defined. Lys146 and Phe132 are comparable. Trp93 shows clear holes for both the five- and six-member rings for both the original data and PASR. For both the original data and PASR processing, the density of tyrosine is dependent on location; Tyr34 is weaker, while Tyr168 is clear and strong with a clear hole in the six-membered ring. Density maps are displayed at 5σ. Even at <2 Å resolution, PASR is competitive in terms of clarity with data collected at the higher magnification in counting mode.

**Figure 6:**
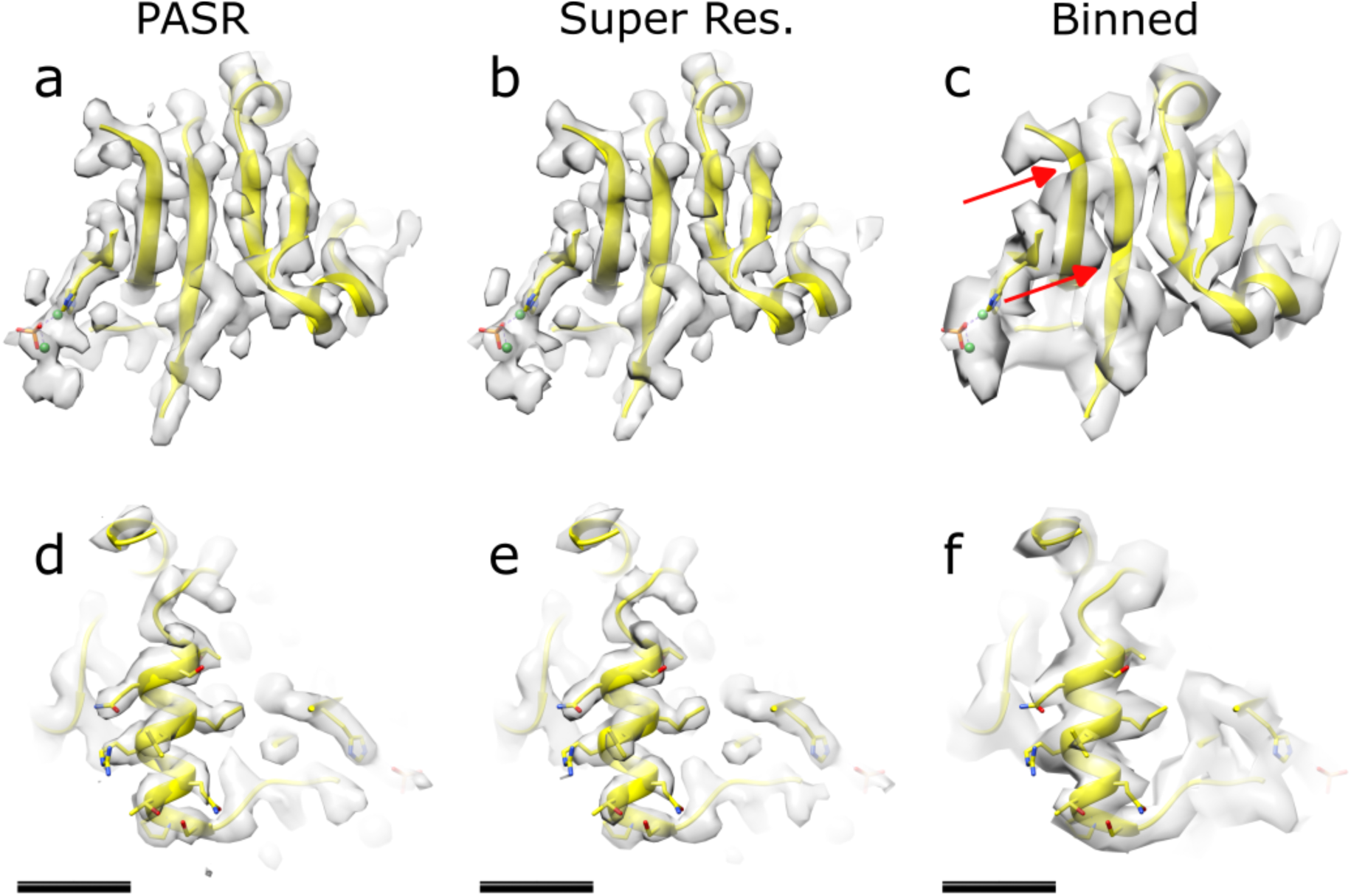
Comparing density of jack bean urease; main chain density of jack bean urease in a β-sheet fold; (a) PASR processed map, (b) super resolution (original data), and (c) limited by physical Nyquist. Note the two main chain density breaks (red arrows) present in the downsampled data, which is not evident for PASR or super resolution, (d) another region of jack bean urease, focussing on an α-helix with sidechains also shown. Clear sidechain density is evident for PASR (d) and super resolution data (e), commensurate with reported resolution. The Nyquist frequency limited data (f) shows ambiguous density for sidechains, particularly the two glutamine residues and the arginine residue. Fitted PDB: 31A4. Once again, here, PASR is clearly competitive with super resolution acquisition, and a large improvement over data limited by physical sampling frequency. Density displayed at 5σ. Scale bar equals 1 nm.

**Figure 7:**
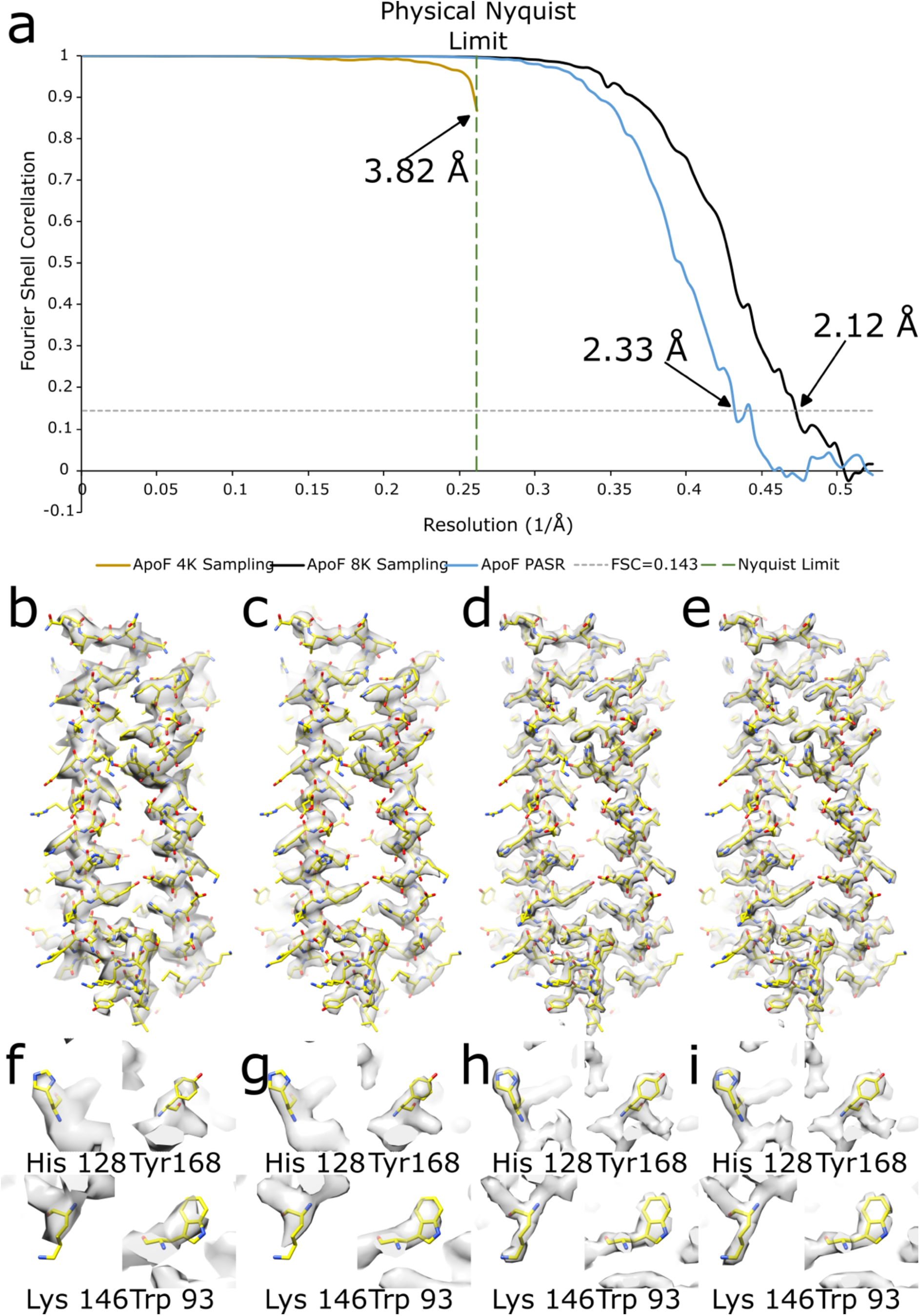
The improvement in resolution with PASR for apoferritin when acquired with conditions which image almost an entire grid hole; (a) FSC curves of the 4K sampled original data (mustard yellow curve), 8K sampled original data (black curve) and PASR processed data (blue curve). FSC=01.43 is indicated with a dashed grey line and the physical Nyquist limit is indicated with a dashed green line. This is a “worst case scenario” for PASR, approximately 0.2 Å lower resolution than “native” super resolution data. The same data processed with CryoSPARC is approximately 0.15 Å lower resolution than the native super resolution data. (b) monomer of apoferritin with monomer rigid-body fitted, (c) the same monomer but resampled to the same pixel spacing as PASR/super resolution data, showing no visible improvement in map clarity, (d) monomer of apoferritin with monomer rigid-body fitted from super resolution (8K sampling) processing, (e) monomer of apoferritin with monomer rigid-body fit from PASR processing. (f-i) example side chain density for each of (b-e). PASR is clearly competitive with the super resolution (8K) sampled data. Density displayed at 5σ.

**Figure 8:**
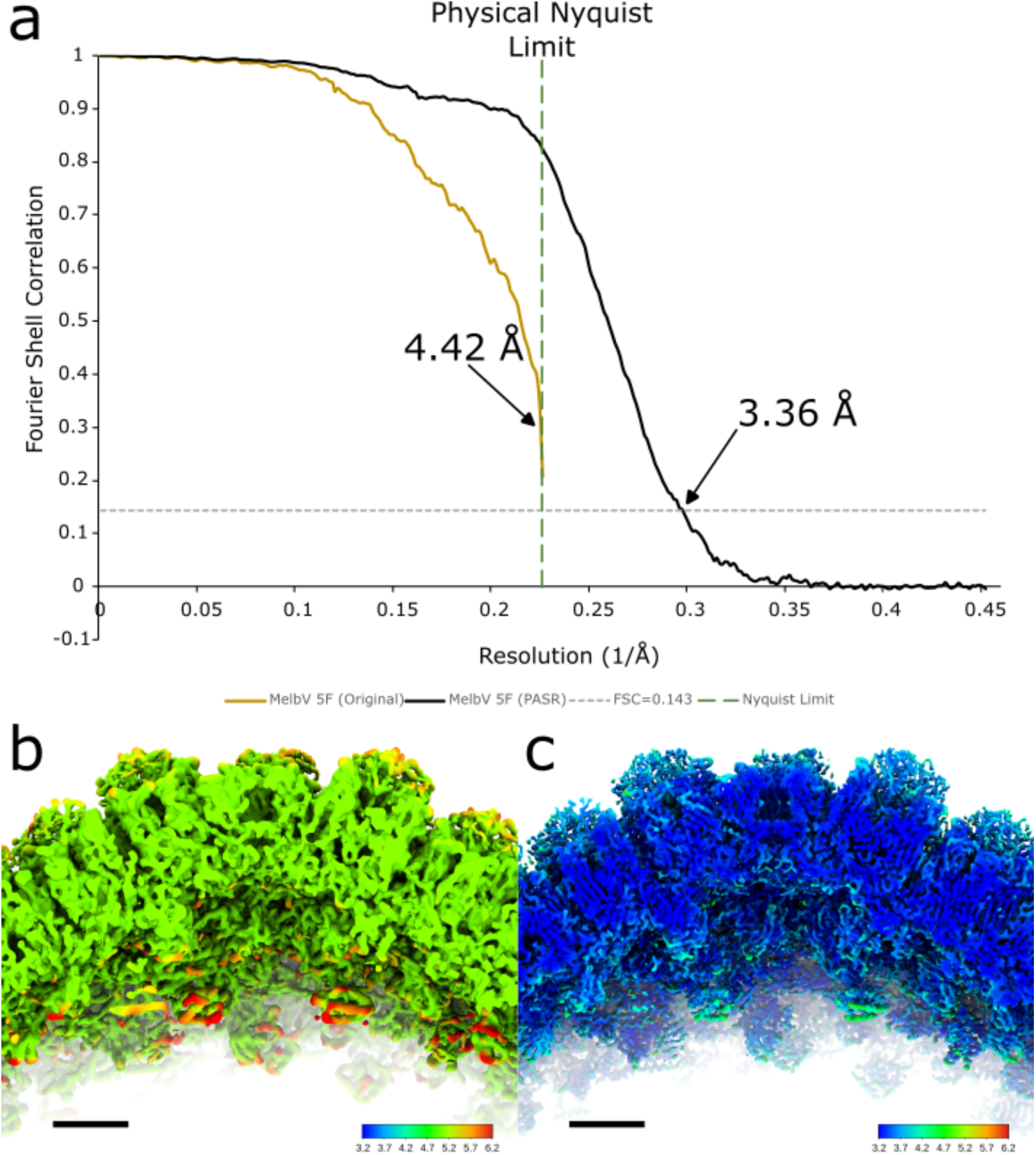
The improvement in resolution with PASR for Melbournevirus, which had originally been limited by sampling frequency during acquisition; (a) gold standard FSC curves of the fivefold block, showing the potential improvement from application of PASR. Original curve (mustard yellow line) and PASR FSC (black line), (b) locally filtered map, coloured by local resolution of a central slice of the original fivefold block, (c) locally filtered map, coloured by local resolution of a central slice of the PASR fivefold block. Maps shown at 3σ. Scale bars equals 5 nm. Colour scales are identical and colour from 3.2 Å (blue) through to 6.2 Å (red). The improvement in main chain clarity is dramatic in the capsid, making model building for proteins of undetermined sequence much more realistic.

**Table 5:** Parameters for Melbournevirus processing.

| Melbournevirus (RELION 3.1) |  |  |
| --- | --- | --- |
|  | Original | PASR |
| Accelerating voltage | 300 | 300 |
| Spherical aberration | 2.7 | 2.7 |
| Micrographs (total) | 3,367 | 3,367 |
| Magnification (Å/pixel) | 2.21 | 1.105 |
| Passed CTF estimation | 3,067 | 3,140 |
| CTF estimation cutoff (Å) | Manual | 10 |
| Particles picked | 7,000 | 7,747 |
| Particles selected | 3,852 | 5,312 |
| Final resolution (5F/3F/2F) (Å) | 4.42/4.42/4.43 | 3.48/3.48/3.36 |
| Final particle count (sym. expanded) | 3,124 (187,440) | 3,919 (235,140) |
| Estimated b-factor | -100 (imposed) | -100 (imposed) |

**Table 6:** Parameters for adeno-associated virus processing.

| Adeno associated virus (RELION 3.1) |  |  |
| --- | --- | --- |
|  | Original | PASR |
| Accelerating voltage | 300 | 300 |
| Spherical aberration | 2.7 | 2.7 |
| Micrographs (total) | 6,067 | 6,067 |
| Magnification (Å/pixel) | 1.34 | 0.67 |
| Passed CTF estimation | 6,001 | 5,995 |
| CTF estimation cutoff (Å) | Manual | Manual |
| Particles picked | 1,473,552 | 1,614,326 |
| Particles selected | 1,090,590 | 1,100,376 |
| Final resolution (Å) | 2.68 | 2.16 |
| Final particle count | 1,090,590 | 1,023,144 |
| Estimated b-factor | -98.8 | -100.8 |

PASR can function on pre-motion-corrected micrographs if the data reaches the Nyquist limit if applied to either the raw micrographs or a particle stack. However, the improvement is markedly less (Fig. S17) than either native data or on PASR applied prior to motion correction (Fig. S1-5). We used Nodavirus from EMPIAR 10203 (Ho et al., 2018) (Table 7) to test further and downsampled the micrographs in Fourier space by a factor of two. This made the effective sampling limit 2.12 Å/pixel. A reconstruction with this binned data reached the sampling limit (Fig. S18). Applying PASR to these downsampled micrographs and then reprocessing permitted recovery of information up to a gold standard FSC of 3.21 Å (Fig. S19). This data was originally published at 3.28 Å (Ho et al., 2018). We have previously reprocessed this data and achieved 2.9 Å (Fig. S20), so this is not the feat it may appear.

**Table 7:** Parameters for Nodavirus (EMPIAR −10203) processing, which is comprised of pre-motion-corrected micrographs, not movies.

| Nodavirus (EMPIAR-10203) (RELION 3.1) |  |  |  |
| --- | --- | --- | --- |
|  | Original | Binned | PASR |
| Accelerating voltage | 300 | 300 | 300 |
| Spherical aberration | 2.7 | 2.7 | 2.7 |
| Micrographs (total) | 2,459 | 2,459 | 2,459 |
| Magnification (Å/pixel) | 1.06 | 2.12 | 1.06 |
| Passed CTF estimation | 2,457 | n/a | n/a |
| CTF estimation cutoff (Å) | Manual | n/a | n/a |
| Particles picked | 66,615 | n/a | n/a |
| Particles selected | 29,180 | Used original particle stack | Used original particle stack |
| Final resolution (Å) | 2.86 | 4.24 | 3.21 |
| Final particle count | 29,180 | n/a | n/a |
| Estimated b-factor | -108.1 | -57 | -143.9 |

### Application tests for PASR

Rather than reprocess everything again in CryoSPARC (Punjani et al., 2017), we tested the Apo64K dataset (Table 2, Fig. S4, S5). This subset was originally collected as part of a much larger dataset (Burton-Smith et al., to be published) on the Titan Krios G4 (TFS) at the National Institute for Physiological Sciences with a Falcon 4 camera. As the Falcon 4 EER format allows sampling at 4K (native), 8K (super resolution) or 16K (double super resolution), we pre-processed TIFF converted micrographs at 4K with PASR and compared it to 8K sampling. The CryoSPARC pipeline (or the basic parts, at least) show no anomalous results with PASR pre-processed data (Fig. S5). All reconstructions resulted in maps which were visually appropriate for the reported resolution with no artefacts visible in the maps, half maps or diagnostic metric output.

Finally, for one last test, we zero-padded the half maps from one of the Melbournevirus reconstructions, rescaled the mask appropriately and carried out post processing (Fig. S21). As is expected with zero padding, the FSC of the unmasked half maps falls to zero at the original sampling limit and flatlines (green curve, Fig. S21a). However, the phase randomised FSC had fallen to zero where randomisation starts but begins to increase again once past the original sampling limit (red curve, Fig. S21a). When RELION was left to estimate b-factor, rather than manually imposing one, the estimated b-factor exceeded −1500. Furthermore, local resolution estimation reports exactly half the local resolution estimation from the non-zero padded data (Fig. S21b). PASR processed data does not demonstrate these artefacts. Finally, the map looks like one which has been rescaled (resampled) after completion (e.g.: in UCSF Chimera (Goddard et al., 2007; Pettersen et al., 2004) or with relion_image_handler) (Fig. S21c).

## Discussion

Here we demonstrate a super-resolution pre-processing step for cryo-EM data that is conventionally limited by Nyquist. All data subjected to PASR pre-processing behaved as one would normally expect when using the popular RELION suite (Fernandez-Leiro & Scheres, 2017; Scheres, 2012). The dataset tested in CryoSPARC (Punjani et al., 2017) also behaved normally. For the dataset requiring initial model generation in cisTEM (Grant et al., 2018) it also behaved normally, although a full processing pipeline was not carried out. As time and resources permit, we will test the other software, and encourage others to do so as well.

While in some datasets fewer micrographs passed CTF estimation when PASR pre-processed, most micrographs were still perfectly acceptable, and defocus, astigmatism and maximum resolution estimated by CTFFIND (Rohou & Grigorieff, 2015) for original and PASR pre-processed micrographs were within acceptable margins of error (Fig. 3). Perhaps the micrographs which failed CTF estimation with PASR pre-processing did so due to decreased contrast, as both super resolution and PASR data demonstrates lower contrast than “native” data, and downsampling native data also decreases micrographs which pass CTF estimation. Or, perhaps they were close to failing and may have done so with a different version of CTFFIND, as we have noted that some releases of CTFFIND appear more forgiving than others for difficult micrographs. CTF estimation worked when using both the motion corrected micrographs for CTF estimation (Fig. S6-S16) and the independent power spectrum (Fig. S1-3) which can be optionally generated by RELION when using the RELION implementation (Zivanov et al., 2018) of the MotionCor2 (Zheng et al., 2017) algorithm. Patch CTF in CryoSPARC (Punjani et al., 2017) worked as expected (Fig. S4, S5).

Previously, the solution to reaching the Nyquist limit would be to collect more data at a higher magnification. However, sometimes this is not possible, or is even undesirable due to the type of sample. The resolution limit in the Melbournevirus dataset (Burton-Smith et al., 2021) was the *raison d’être* for this investigation. Acquiring giant virus datasets at higher magnifications can lead to tens or hundreds of thousands of micrographs for the same, or fewer, usable particles (Wang et al., 2019), with the possibility that the data will not achieve a resolution commensurate with that magnification because of the myriad other factors which influence cryo-EM SPA, and particularly giant virus reconstructions. Our Tokyovirus data (Chihara et al., 2022) were collected with super resolution, which dramatically slowed micrograph acquisition, but we did not reach what would have been the sampling limit of counting mode (Fig. 1b) acquisition. Conversely, in our Melbournevirus report (Burton-Smith et al., 2021) data were collected in counting mode, but reached the physical sampling limit. Size is not the only challenge of processing giant viruses. For example, they can demonstrate pseudosymmetry – the block-based reconstruction technique was applied to PBCV-1 with icosahedral symmetry initially (Fang et al., 2019), however it is not icosahedral in form (Shao et al., 2022). Likewise, Mimivirus was initially analysed as icosahedral in form (Xiao et al., 2005), but has a “stargate” which means it demonstrates only fivefold symmetry (Xiao et al., 2009). Further, the capsids can be frustratingly heterogeneous (Watanabe et al., 2022), demonstrating flexibility which limits achievable resolution even at lower magnifications. Then, perhaps, it would be better to collect data at a lower magnification and increase the probability of reaching resolutions where PASR might be applied to exceed that sampling limit, if necessary. For working with giant viruses, flexibility in both acquisition and processing are critical. PASR processing of the Melbournevirus data yielded maps from block-based reconstruction (Zhu et al., 2018) which already exceeded the original sampling limit, when Bayesian polishing was applied (using the default sigma parameters) the final gold standard FSC reported for each block was 3.5 Å for the twofold block (Fig. S6), 3.5 Å for the threefold block (Fig. S7) and 3.4 Å for the fivefold block (Fig. 8, S8) respectively, with local resolution extending to ~3.2 Å for all three blocks (Figs. S6-S8).

Other datasets (Table 1) were used to demonstrate that PASR is applicable to several complexes acquired with some of the most common detectors (Falcon 3, Falcon 4, K2 Summit, K3) and microscopes (200 and 300 kV). The downsampling (Fig. S15) and PASR (Fig. 5, S14) reprocessing of a previous public apoferritin dataset (Fig. 5, S16) was used so we could directly compare to “native” data. The downsampling processing of individual frames was necessary so that PASR could be applied to a “Nyquist limited” micrograph movie dataset, as (at time of writing) except for EMPIAR-10549 (Feathers et al., 2021), there are no public datasets which exceed the physical Nyquist limit (however, EMPIAR-10025 (Campbell et al., 2015) does get quite close with modern processing suites).

Similarly, the downsampling (Fig. 7, S1) of the super resolution (Fig. 7, S2) dataset before PASR (Fig. 7, S3) pre-processing of the downsampled data was used to permit a comparison to “native” super resolution, i.e., super resolution data acquisition time. Acquisition-time super resolution results in a slightly superior resolution compared to PASR as the sub-pixel sampling means that of the 2×2 square which corresponds to a physical pixel on the camera, the output pixels are unique, while PASR means they are identical. With PASR, the motion correction step allows the motion of the frames to treat each sub-pixel independently, resulting in unique pixels in the motion corrected micrographs. The same is true for per-particle alignment in Bayesian polishing.

The AAV dataset (Fig. S9) was used as another test (after the Melbournevirus dataset) because it was acquired with a different microscope and detector. AAV is also much smaller than Melbournevirus and thus requires no block-based reconstruction (Zhu et al., 2018) to process. The AAV dataset was acquired with a moderately high defocus, however, which may have created a limit in the achievable resolution, nevertheless, it still exceeded the original sampling limit (Fig. S10) although we had been aiming for <2 Å. High symmetry means increased likelihood in achieving close to the Nyquist limit as symmetry acts as a particle multiplier. The urease dataset (Fig. 6, S11) was chosen as it is the only public dataset which exceeds physical Nyquist; however, it had the second advantage of being (relatively) low symmetry, thus permitting us to examine the effects of PASR on lower symmetry samples, where it was still shown to provide benefits (Fig. 6, S12). The downsampled data provided a benchmark for how non-super resolution data would perform (Fig. S13) and curiously demonstrated some breaks in the main chain (Fig. 6c) of the structure which neither the super resolution nor the PASR data demonstrated (Fig. 6a, b).

PASR does *not* exhibit the issues which “zero padded” half-maps demonstrate on the FSC or local resolution estimation. Zero padding half maps before FSC curve calculation results in no improvement in the FSC (green curve, Fig. S21a) as the values are simply zero. However, zero padding does exhibit a clear effect on the phase randomised FSC (red curve, Fig. S21a), the masked map FSC (blue curve, Fig. S21a) and corrected FSC (black curve, Fig S21a) which should make it immediately obvious something is wrong. An estimated b-factor in excess of −1500 from RELION also means the output map is a ball of noise. With zero padded half maps, local resolution estimation estimated 2.2 Å for the capsid of the Melbournevirus block (Fig. S21b), but the FSC is reported as 4.42 Å (the original sampling limit) and visual examination of the map looks like 4.4 Å (Fig. S21c).

While initial model generation in the PASR-processed jack bean urease (Fig. S12) dataset failed using the RELION 3.1 stochastic gradient descent algorithm, it succeeded using cisTEM (Grant et al., 2018). As such, given the challenges of initial model generation we do not consider this to be a consequence of PASR processing. Further, it would be advisable to apply PASR only to a dataset which has already achieved Nyquist resolution. This would mean the dataset already has a good initial model from the original data, and that *ab initio* model generation is not a requisite of PASR pre-processed data.

PASR works to improve the resolution and clarity of cryo-EM datasets which are Nyquist limited but does not “magically” provide extra resolution. Unless you have already achieved a reconstruction which is limited by the Nyquist frequency, it will not help. The improvement in FSC below the physical Nyquist limit for the PASR processed Melbournevirus dataset (Fig. 8) was likely due to application of Bayesian polishing, although the reconstructions exceeded the original sampling limit before applying Bayesian polishing. Further, as box sizes double, it will slow processing commensurately, just as it will for acquisition time super resolution data.

PASR does not currently compete “neck-and-neck” with data collected natively at the equivalent higher magnification, and likely never will, however the improvement over non-super resolution data is clear (Figs. 5-8, S1-16). While reporting gold standard FSCs around 0.07-0.2 Å lower than native super resolution data, side chain clarity is comparable, which does make us question how valuable the reported (global) resolution is as a metric of performance. In effect, there are definitive “bands” across the resolution range (Rosenthal & Rubinstein, 2015) at which point features are lost or gained, but within that band minor changes in reported resolution mean little (Fig. 5-7). B-factor is also marginally degraded (Tables 2–8), although the impact of this is debatable as there is no hard and fast rule regarding what sharpening should be applied to a map, and it will often be manually adjusted for clarity. Those aiming for record-breaking resolutions should still collect data at higher magnification. Further, from the urease (Figs. 6, S11-13) and Apo64K (Figs 7, S1-5) datasets, PASR does not quite achieve parity with acquisition-time super resolution. This is not entirely unexpected, as acquisition-time super resolution treats each sub-pixel independently and each sub-pixel in each frame possesses a unique value. PASR, however, is limited in that the sub-pixels must be treated as identical, to become unique after motion correction. On the other hand, it also occupies one quarter of the storage space, assuming they are stored in comparable formats.

**Table 8:** Parameters for apoferritin processing for small dataset at low magnification using CryoSPARC.

| Apoferritin (this study) (CryoSPARC) |  |  |
| --- | --- | --- |
|  | Original (8K) | PASR |
| Accelerating voltage | 300 | 300 |
| Spherical aberration | 2.7 | 2.7 |
| Micrographs (total) | 32 | 32 |
| Magnification (Å/pixel) | 0.956 | 0.956 |
| Passed CTF estimation | 32 | 32 |
| CTF estimation cutoff (Å) | n/a | n/a |
| Particles picked | 57,234 | 57,988 |
| Particles selected | 47,695 | 45,783 |
| Final resolution (Å) | 2.28 | 2.45 |
| Final particle count | 47,695 | 45,783 |
| Estimated b-factor | -74.8 | -114.4 |

Collecting at a lower magnification means more particles per micrograph, which will play a significant role in increasing particle count for giant virus cryo-EM SPA, where there can be as little as a single particle per micrograph (Wang et al., 2019). Likewise, storage space is decreased as it is not necessary to record and archive raw super resolution micrograph movies. As these are approximately four times the size of counting mode micrographs (when using the same storage format) this should ease the burden on cryo-EM facilities data storage and archival. Equally, the validations carried out with public datasets required pre-processing the micrographs before PASR treatment, which may have caused loss (by comparison to the Melbournevirus dataset) of resolution. However, the direct comparison of PASR to 8K sampled micrographs from a Falcon 4 do show performance to be slightly behind that of acquisition time super resolution. What is chosen will depend on the equipment available to the facility, nature of the sample and ultimate objective of the study.

Currently, the PASR pre-processing step does take some time. Particularly, there is a significant bottleneck in reading and writing TIFF stack files using GNU Octave, although incorporating this method into a native program would solve this, as writing TIFF files is much faster in, e.g., the *mrc2tif* program of IMOD (Kremer et al., 1996). The Octave version permitted step-by-step validation that the PASR pre-processing was happening as intended. A Python implementation is complete, which can be applied much more quickly. We think it would be best to implement PASR into the processing suite pipelines natively, such that it could be applied in volatile memory during motion correction. Alternatively, rather than be required to completely reprocess a dataset, it may be better to implement PASR at a later stage in the processing pipeline, perhaps as an option akin to Bayesian polishing (Zivanov et al., 2019) in RELION, which is only carried out after early processing and reconstructions are complete. This can be viewed in the same way as downsampling during early processing. Micrograph movies could be rescaled in volatile memory, motion correction and/or polishing performed, and particles re-extracted without requiring the (temporary) storage of a pre-processed dataset four times the size of the original. This in turn means larger datasets may be collected. The current implementation of PASR is to allow manual validation on a per-frame basis that output is as expected.

One thing we have not tested is whether PASR can be utilised on data already acquired with acquisition-time super resolution at lower magnifications resulting in reaching the Nyquist limit in the (non-downsampled) super resolution data. This may be examined in the future, however the low DQE of super resolution data at the highest frequencies will almost certainly mean that reaching “super resolution Nyquist” – the frequency at which PASR would become valuable – is difficult. In fact, our recent work (Burton-Smith et al. to be published) demonstrates that while it is possible to reach and exceed super resolution Nyquist, it is something of a struggle to do so. We briefly tested RELION 4 and the VDAM algorithm (Figs. 7, S6-8), along with CryoSPARC (Figs. S1-2) on PASR data with comparable results to RELION 3.1 (Figs. 5,6,8, S3-5, S14-21).

PASR permits a super resolution-like exceeding of the Nyquist limit on datasets which have proven to reach that limit. Our experiments with public data show that it does not perform quite as well, in terms of final reported resolution, as data which has been acquired natively at the equivalent magnification (Fig. S3, S5). For data limited by Nyquist, such as our recent Melbournevirus report (Burton-Smith et al., 2021), PASR provides a method for exceeding this limit and improving both the resolution and clarity of reconstructions (Fig. 8, S14-16). Using a modern model building suite, Model Angelo (Jamali et al., 2023), we could build candidate protein structures into the Melbournevirus capsid with some success (Burton-Smith et al., to be published). Comparatively, model building into the 4.4 Å map failed completely for Model Angelo, while DeepTracer (Pfab et al., 2021) struggled and eventually the MCP (major capsid protein) was built by hand (Burton-Smith et al., 2021).

While we were preparing this manuscript, Cryo-ZSSR (Huang et al., 2023) was brought to our attention. Cryo-ZSSR follows a similar concept to PASR but utilises a neural network implementation, and examining the documentation appears to have several other limitations not evident with PASR. The movies require complex pre-processing to generate intermediate movies; for each intermediate movie frame generated the whole cryo-ZSSR pre-processing step must be repeated, rather than the simpler pixel doubling implementation of PASR followed by a standard cryo-EM processing pipeline. In large datasets containing micrographs with many frames this will be slower than either the current PASR implementation or our aim of PASR purely in volatile memory. Further, as they admit, the current implementation of cryo-ZSSR will automatically crop input to 512×512 pixels – with detectors outputting 4,096×4,096 (Falcon 3, 4), 3,838×3,710 (K2 Summit) or 5,760×4,092 (K3) this appears a severe limitation to the current implementation. In our experiments, PASR reports resolutions closer to the original data than cryo-ZSSR appears to, although we are intrigued by their method.

In conclusion, we envision this technique to be of greatest utility to cryo-EM researchers working on giant structures where the number of objects which can be imaged on each micrograph is limited but hope others may find the technique of use. Large cryo-EM facilities face ever increasing storage demands for raw data so may find PASR attractive for samples known to be heterogeneous. Possibly, *in vivo* tomography and sub-tomogram averaging may also find PASR of utility: more potentially “good” particles can be acquired per tomogram due to increasing the observed area of each lamella, perhaps paired with something akin to PACE-Tomo (Eisenstein et al., 2022). This increases the likelihood of acquiring high quality averages of the subject of interest or capturing more possible states of a complex. Facilities which do not have detectors capable of super resolution acquisition may also find PASR of interest – the highest resolution cryo-EM reconstruction published at time of writing is from data collected from a Falcon 3 (Yip et al., 2020) – so detectors without super resolution acquisition are still extremely capable. We are working on the “on the fly” PASR implementation so that it may be applied during late-stage processing in a similar manner to Bayesian polishing (RELION) (Zivanov et al., 2019) or per-particle local motion refinement (CryoSPARC) (Punjani et al., 2017), which will be more efficient for storage and processing and is the ultimate target for the full implementation of PASR.

## Materials and Methods

### Datasets used

Table 1 details the datasets used. We chose to test a variety of data based on the use of different detectors for acquisition. EMPIAR-10216 (Danev et al., 2019) was used for apoferritin, as it was acquired using a Falcon 3 direct detector, EMPIAR-10549 (Feathers et al., 2021) was used to test both a direct comparison to K3 super resolution data and a lower symmetry complex. EMPIAR 10203 (Ho et al., 2018) was used for testing the efficacy of PASR when applied to single frame K2 micrographs. We were granted permission to test the adeno-associated virus dataset by Prof. Uchiyama (University of Osaka) as it was collected from a JEOL CRYOARM 300 microscope using a Gatan K3 in CDS (correlative double sampling) mode. Melbournevirus was previously processed (Burton-Smith et al., 2021). The first 32 micrographs of an apoferritin dataset acquired with a Falcon 4 DED used in Burton-Smith *et al*. (to be published) was tested as it allowed us to test both “native” (4K) and super resolution (8K) processing from a single dataset without requiring binning. Datasets not currently public, and the PASR preprocessed micrographs will be made available as soon as possible via EMPIAR (Iudin et al., 2016). All datasets were processed with the “Ignore CTFs until first peak?” parameter set to “Yes” in the RELION user interface.

### Pre-processing

A script (see Supplementary Information) was written in GNU Octave (Eaton et al., 2019; Eaton et al., 2022) supplemented by the Image package (https://gnu-octave.github.io/packages/image/), and the ReadMRC, WriteMRC and WriteMRCHeader functions developed for MATLAB (The MathWorks, 2022) by F. Sigworth (Sigworth, 2023) to import micrograph movies, divide each pixel into a 2×2 grid with each sub-pixel having an identical value to the originating pixel, and write the resulting “super resolution” micrograph movie to a different file. A modified form, using the ReadTIFFStack function, was used for TIFF format micrographs. This script carried out “pixel doubling” on each frame of every micrograph movie as described above. These pre-processed movies were passed to the standard RELION (Fernandez-Leiro & Scheres, 2017) processing pipeline. RELION 3.1 was used except for cases explicitly stating RELION 4 or CryoSPARC. To test EMPIAR-10216 and -10549 with PASR, another script was written which first binned down each 2×2 area to a single pixel by Fourier space cropping. These downsampled micrograph movies were then reprocessed with the PASR script.

### Processing of apoferritin dataset (Apo64K) from NIPS Titan Krios G4

This data was processed in RELION 4. Table 3 summarises this. The 4K sampled original data were imported and motion corrected with the RELION (Zivanov et al., 2018) implementation of the MotionCor2 algorithm (Zheng et al., 2017) with 32-bit MRC output and separate power spectra. Separate power spectra and the micrographs reported highly similar CTF value estimates with CTFFIND (Rohou & Grigorieff, 2015). A small number of particles were manually picked and classified, before template picking was used to select a total of 102,364 particles. After 2D classification, 33,468 particles were selected in clear classes and an initial model generated. After a single round of 3D refinement, the reconstruction reached the physical Nyquist limit.

The 8K sampled original data were imported and motion corrected with the RELION (Zivanov et al., 2018) implementation of the MotionCor2 algorithm (Zheng et al., 2017) with 32-bit MRC output and separate power spectra. Separate power spectra and the micrographs reported highly similar CTF value estimates with CTFFIND (Rohou & Grigorieff, 2015). A small number of particles were manually picked and classified, before template picking was used to select a total of 84,586 particles. After 2D classification, 63,313 particles were selected in clear classes and an initial model generated. After cycling 3D refinement and CTF refinement (magnification anisotropy, beam tilt, per-particle defocus, and astigmatism), followed by Bayesian polishing, the final gold standard FSC reported 2.13 Å with a b-factor of −62.4.

The PASR processed data were imported and motion corrected with the RELION (Zivanov et al., 2018) implementation of the MotionCor2 algorithm (Zheng et al., 2017) with 32-bit MRC output and separate power spectra. Separate power spectra and the micrographs reported highly similar CTF value estimates with CTFFIND (Rohou & Grigorieff, 2015). A small number of particles were manually picked and classified, before template picking was used to select a total of 102,493 particles. After 2D classification, 64,597 particles were selected in clear classes and an initial model generated. After cycling 3D refinement and CTF refinement (magnification anisotropy, beam tilt, per-particle defocus and astigmatism), followed by Bayesian polishing, the final gold standard FSC reported 2.33 Å with a b-factor of −99.

The 8K sampling original data was imported into CryoSPARC (4.2.1) (Punjani et al., 2017) and patch motion correction and patch CTF estimation were carried out with the default parameters. Blob picking was used, selecting a ring blob of internal diameter 90 Å and external diameter 110 Å, which selected 57,234 particles. Particles were extracted and 2D classified into 100 classes with 250 particles per class and a maximum resolution of 4 Å which seems to distinguish apoferritin classes more clearly. The clear classes, containing 47,695 particles were passed to *ab initio* model generation and homogeneous refinement with defocus and beam tilt optimisation enabled, resulting in a final resolution of 2.28 Å with a b-factor of −74.8.

The PASR processed data was imported into CryoSPARC (4.2.1) (Punjani et al., 2017) and patch motion correction and patch CTF estimation were carried out with the default parameters. Blob picking was used, selecting a ring blob of internal diameter 90 Å and external diameter 110 Å, which selected 57,988 particles. Particles were extracted and 2D classified into 100 classes with 250 particles per class and a maximum resolution of 4 Å which seems to distinguish apoferritin classes more clearly. The clear classes, containing 45,783 particles were passed to *ab initio* model generation and homogeneous refinement with defocus and beam tilt optimisation enabled, resulting in a final resolution of 2.45 Å with a b-factor of −114.8.

### Processing of apoferritin dataset (EMPIAR-10216)

The original data, the downsampled data and the PASR data were processed via normal RELION processing. Table 3 summarises this. The original dataset was processed by importing all micrographs into RELION 3.1 and following a standard processing pipeline of motion correction, CTF estimation, of which 1,228 micrographs passed CTF estimation with a resolution <5 Å. Template based particle picking selected 217,495 particles, of which 161,966 were selected in clear classes after 2D classification. 3D classification was carried out for initial particle alignment before 3D refinement. Parameter optimisation with CTF refinement and Bayesian particle polishing were carried out before Ewald sphere curvature correction was applied to the final reconstruction to a final resolution of 1.7 Å with an estimated b-factor of −44.4.

The downsampled dataset was processed in the same way, with 1,153 micrographs passing CTF estimation with a 5Å cutoff, with template autopicking selecting 156,099 particles. 149,536 particles were selected in clear classes after 2D classification, before 3D classification and refinement. After post processing this refinement reached the downsampled sampling limit before any CTF refinement, polishing or Ewald sphere curvature correction was required, so they were not applied. The final resolution was 2.068 Å with an estimated b-factor of −46.8.

The PASR dataset was also processed in the same way. After motion correction and CTF estimation, 1,106 micrographs had an estimated CTF resolution of <5 Å. Template autopicking selected 207,388 particles. 152,367 particles were selected in clear classes, although in the 2D classification step, the number of clear classes was significantly reduced compared to both the original and binned data. After CTF refinement (magnification anisotropy, beam tilt and particle defocus/astigmatism refinement), Bayesian polishing and Ewald sphere curvature correction, the final resolution was 1.78 Å with an estimated b-factor of −50.2.

### Processing of jackbean urease dataset (EMPIAR-10549)

The original data, the downsampled data and the PASR data were processed via normal RELION processing. Table 4 summarises this.

The original dataset was processed by importing 286 micrographs and motion correcting with MotionCor2. After CTF estimation, 281 micrographs demonstrated good fits. Laplacian-of-Gaussian (LoG) particle picking was used with a minimum diameter of 110 Å and a maximum diameter of 130 Å. 1,175,198 particles were extracted and 2D classified with clear classes selected containing 837,778 particles. An initial model was generated with D3 symmetry, which was used as a reference for initial alignment with 3D classification before 3D refinement. Parameter refinement was carried out to optimise per-particle defocus and astigmatism, magnification anisotropy and beam tilt. Bayesian polishing was carried out before post processing estimated a final resolution of 3.07 Å with an estimated b-factor of −156.7.

The downsampled dataset was processed by importing the micrographs and motion corrected with MotionCor2 (Zheng et al., 2017). After CTF estimation, 279 micrographs passed. LoG picking was used, with a minimum diameter of 110 Å and a maximum diameter of 130 Å, for a total of 1,170,221 particles, of which 747,661 were within clear classes after 2D classification. An initial model was generated and used as an initial reference in 3D classification. After a single 3D refinement, the physical Nyquist limit was reached of 4.2 Å with an estimated b-factor of −218.8.

The PASR dataset was processed by importing the 286 micrographs and motion correcting with MotionCor2 (Zheng et al., 2017). After CTF estimation, 280 micrographs were passed to LoG autopicking, with a minimum diameter of 110 Å and a maximum diameter of 130 Å, for a total of 1,173,581 particles. After 2D classification, 740,059 particles were contained within clear classes. Initial model generation in RELION failed twice, so the stack was exported using relion_stack_create and cisTEM (Grant et al., 2018) was used which successfully generated an initial model. This initial model was used for particle alignment with a single 3D classification before 3D refinement was carried out. Parameter optimisation with CTF refinement (magnification anisotropy, beam tilt and per-particle defocus and astigmatism) were carried out followed by Bayesian polishing resulting in a final map of 3.19 Å with an estimated b-factor of −178.4.

### Processing of Melbournevirus

The original dataset was not reprocessed. Please see Burton-Smith et al. (2021) for further details of the original processing. The PASR dataset was processed in a manner as close to the original dataset as possible, however particle selection was Laplacian-of-Gaussian picking only, followed by manual removal of particles at the edges of micrographs before 2D classification. This resulted in ~800 more particles in the whole virus consensus reconstruction compared to our original work. Table 5 summarises this. As the dataset was collected originally at two different occasions and the two datasets contain different numbers of frames, when Bayesian polishing was carried out, it was first carried out on the first dataset, then the second with the same parameters (RELION defaults) and the two sets of polished particles recombined for each focussed block of the virus. The twofold block was processed with a box size of 800 pixels, the threefold block was processed with a box size of 900 pixels and the fivefold block was processed with a box size of 660 pixels. These were chosen as a good balance of coverage of the viral capsid for volume size and processing time, while allowing full coverage of the capsid.

### Processing of adeno-associated virus (AAV)

The original data and the PASR data were processed normally in RELION. Table 6 summarises this. The original dataset was processed by importing all 6,067 micrographs and motion correcting with MotionCor2 (Zheng et al., 2017). CTF estimation was carried out with CTFFIND 4.1 (Rohou & Grigorieff, 2015) and particles from the 6,001 micrographs which passed CTF estimation were picked with the Laplacian-of-Gaussian (LoG) autopicker (Zivanov et al., 2018), with a minimum diameter of 230 Å and a maximum of 290 Å, resulting in a total of 1,473,552 picked particles. These particles were extracted and 2D classified into 30 classes, of which clear classes were selected containing a total of 1,090,590 particles. An initial model was generated from these particles and a 3D refinement carried out which reached the maximum sampling frequency of 2.68 Å with an estimated b-factor of −98.8.

The PASR dataset was processed in the same way, with the following exceptions: we tested a second round of picking with a template picker, and imposed symmetry was I3 rather than I1. This was because we wanted to test block-based reconstruction (Zhu et al., 2018) with the smaller particle. However, block-based reconstruction of AAV was abandoned as too time-intensive. 5,995 micrographs passed CTF estimation with the same resolution cutoff, were again autopicked using the LoG autopicker with the same parameters, extracted and 2× downsampled before 2D classification into 30 classes. We tested template autopicking with three varied averages which resulted in better centred picks and a total of 1,614,326 selected. After 2D classification into 100 classes, 14 clear classes were selected containing a total of 1,100,376 particles. From these, an I3 symmetry initial model was generated, and a 3D refinement carried out. Particles were re-extracted unbinned and the refined map rescaled to act as a reference structure. CtfRefinement (magnification anisotropy, beam tilt and defocus/astigmatism) was carried out before Bayesian polishing. The final map reached a gold standard FSC resolution estimate of 2.17 Å with a b-factor of −100.8.

### Processing of Nodavirus

Nodavirus micrographs from EMPIAR 10203 (Ho et al., 2018) were imported into RELION and CTF estimated with CTFFIND 4 (Rohou & Grigorieff, 2015). Particles were picked with the LoG autopicker with an inner diameter of 230 Å and an outer diameter of 290 Å. There was a total of 66,615 picks. These were extracted and 2D classified, with clear classes containing 29,180 particles. An initial model was generated and a cycles of 3D refinement and CTF refinement carried out until a final resolution of 2.86 Å and b-factor of −108.1.

The micrographs were binned, and CTF estimated with CTFFIND 4 (Rohou & Grigorieff, 2015). The particle locations from the final output of the original data were rescaled to the smaller micrographs and the particles re-extracted. A single 3D refinement reached the binned Nyquist limit of 4.24 Å with an estimated b-factor of −57.

The binned micrographs were PASR processed and CTF estimated using CTFFIND 4 (Rohou & Grigorieff, 2015). The particle locations from the final output of the original data were used and after a 3D refinement and single defocus optimisation pass, the highest resolution achieved was 3.21 Å with a b-factor of −143.9.

### Visualisation

Viewing of 2D classes, etc. were carried out within either RELION (Kimanius et al., 2021; Scheres, 2012; Zivanov et al., 2020) or CryoSPARC (Punjani et al., 2017) respectively. 3D volumes were visualised in UCSF Chimera (Goddard et al., 2007; Pettersen et al., 2004) and UCSF ChimeraX (Goddard et al., 2018). SEGGER (Pintilie & Chiu, 2012) was used to segment maps.

## Acknowledgements

We thank K. Okamoto for sharing the original Melbournevirus dataset and S. Uchiyama for his generous permission to test PASR on his AAV data; R.N.B.S. thanks A. de Marco, R. Danev, T. Moriya and M. Wolf for valuable discussions, and A. Noble for detailed feedback and help with Python. This work was supported by AMED BINDS under Grant Number JP22ama121005j0001, JSPS KAKENHI Grant Number JP22H04926 and a NIPS grant of data science 2022.

## Data availability

Post processed maps, half maps and FSC curves for each reconstruction is available from the EMDB under the following accession codes:

EMPIAR-10216 (reprocessing): t.b.c (to be confirmed)

EMPIAR-10216 (binned): t.b.c

EMPIAR-10216 (PASR): t.b.c

EMPIAR-10549 (reprocessing): t.b.c

EMPIAR-10549 binned): t.b.c

EMPIAR-10549 (PASR): t.b.c

EMPIAR-10203 (reprocessing): t.b.c

EMPIAR-10203 (binned): t.b.c

EMPIAR-10203 (PASR): t.b.c

Apoferritin (64,000×) (1.91 Å/pixel) 4K: t.b.c

Apoferritin (64,000×) (0.956 Å/pixel) 8K: t.b.c

Apoferritin (64,000×) (0.956 Å/pixel) PASR: t.b.c

Apoferritin (64,000×) (0.956 Å/pixel) 8K (CryoSPARC): t.b.c

Apoferritin (64,000×) (0.956 Å/pixel) PASR (CryoSPARC): t.b.c

Adeno-associated virus (original): t.b.c

Adeno-associated virus (PASR): t.b.c

Nodavirus (reprocessing): t.b.c

Nodavirus (binned): t.b.c

Nodavirus (PASR): t.b.c

Melbournevirus twofold block PASR: t.b.c

Melbournevirus threefold block PASR: t.b.c

Melbournevirus fivefold block PASR: t.b.c

The following post processed maps, half maps and FSC are already deposited on the EMDB under the following accession codes:

Melbournevirus twofold block (original): 31531

Melbournevirus threefold block (original): 31530

Melbournevirus fivefold block (original): 31529

The following datasets are already publicly available on the EMPIAR database: 10203, 10216, 10549.

The following datasets will be made publicly available on the EMPIAR database as soon as possible: Apoferritin (64,000×) (4K, 8K, PASR), Melbournevirus (original, PASR).

