## Supplementary Figures for "Post-acquisition super resolution for cryo-electron microscopy"

Much of the supplementary information are validation runs carried out to perform checks on PASR. Processing summaries for all full runs are included. FSC curves for tests of PASR on pre-motion-corrected micrographs are included, with the maps deposited in the EMDB. Raw data will be uploaded to EMPIAR.

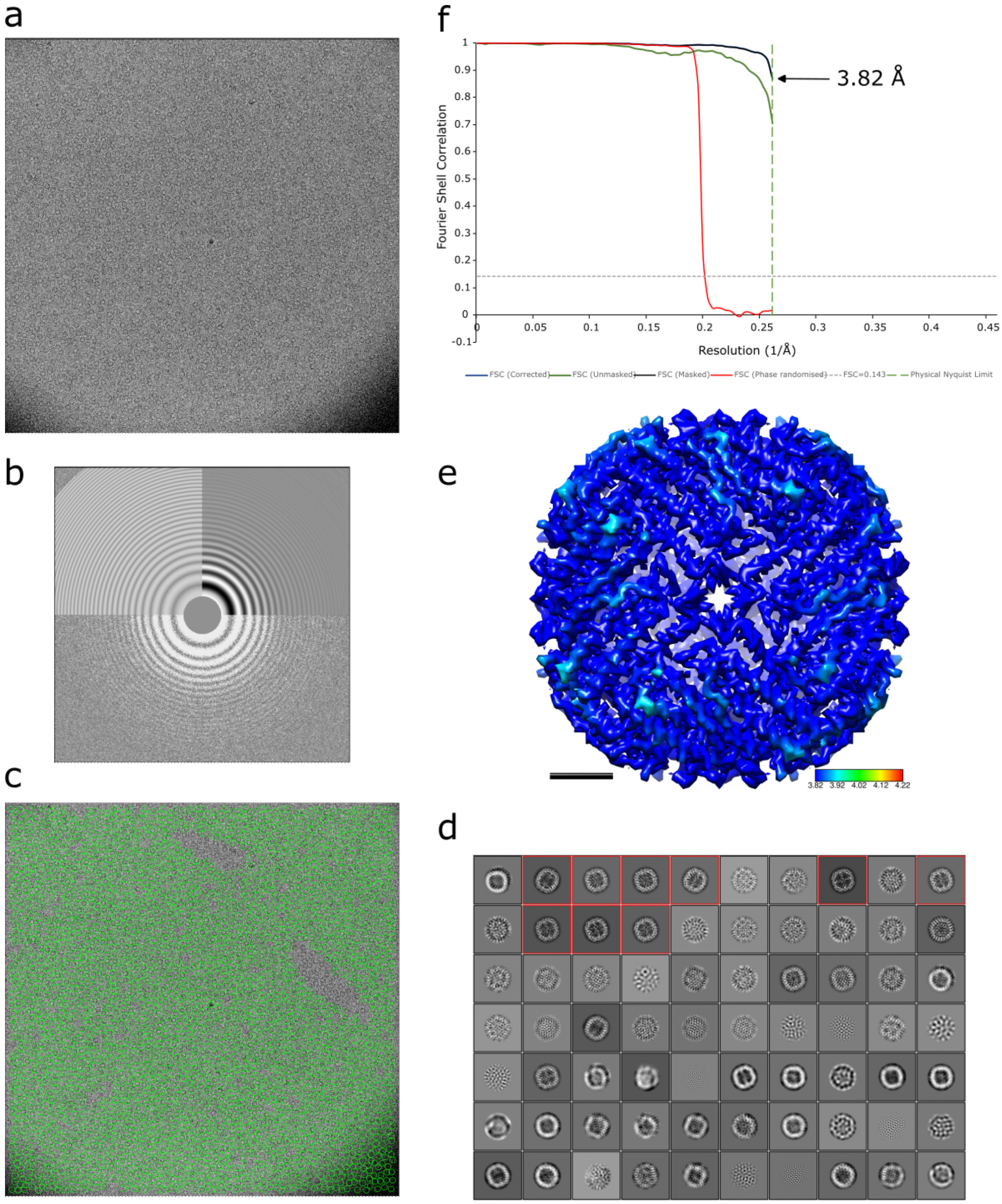


**Figure S1**: Processing of 4K sampled data from second apoferritin dataset. This dataset permits comparison of behaviour with respect to PASR processing for Falcon 4 detectors. (a) Representative micrograph, (b) power spectrum of (a), (c) template picks from (a) are circled in green, (d) result of 2D classification, (e) final reconstruction, coloured by local resolution, with the entire reconstruction estimated at the binned Nyquist limit (3.82 Å) from a global gold standard FSC of 3.82 Å. Scale bar 2 nm. Density displayed at 5σ.

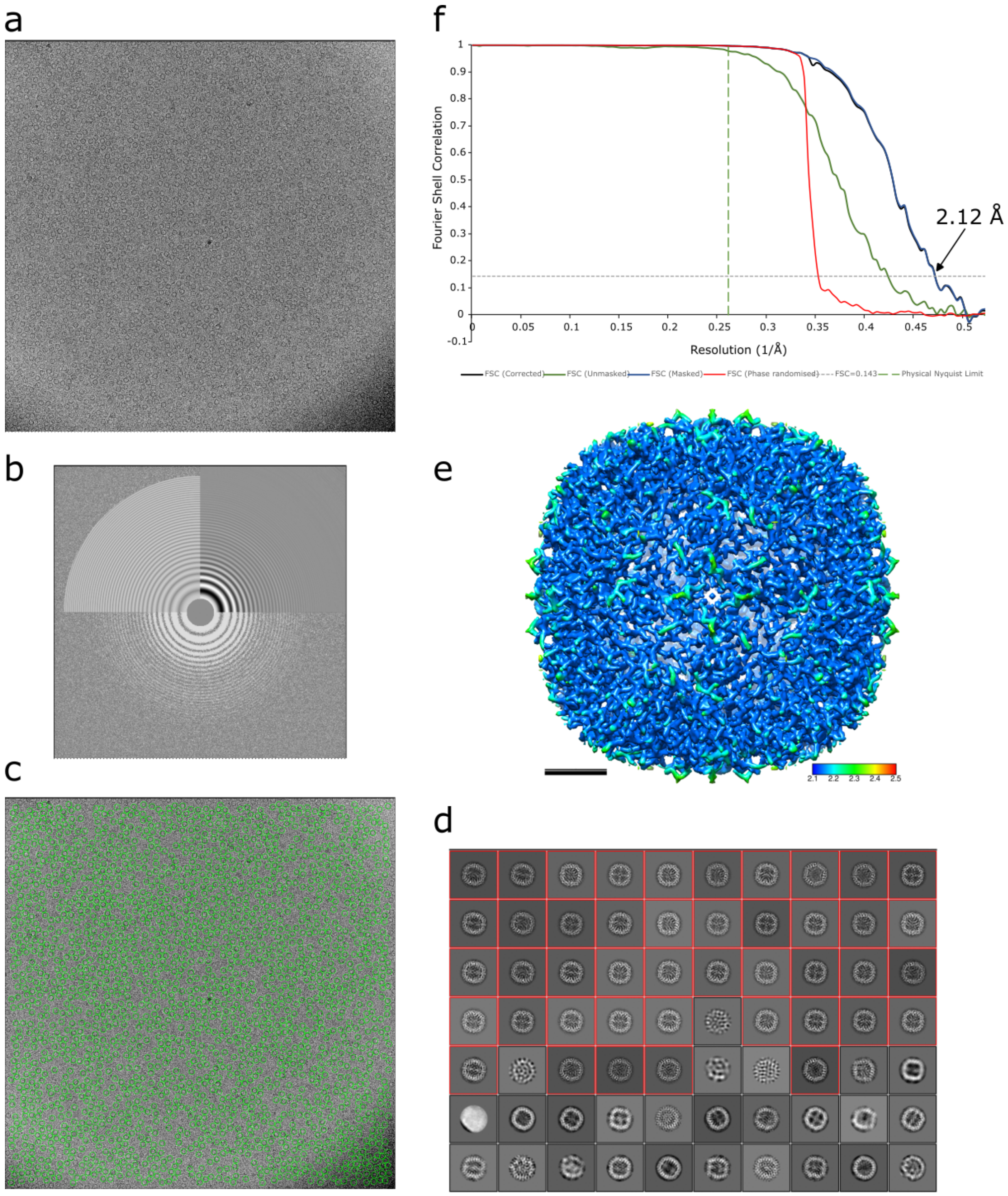


**Figure S2**: Processing of 8K sampled data from second apoferritin dataset. This dataset permits comparison of behaviour with respect to PASR processing for Falcon 4 detectors. (a) Representative micrograph, (b) power spectrum of (a), (c) template picks from (a) are circled in green, (d) result of 2D classification, (e) final reconstruction, coloured by local resolution, extending to a local maximum resolution of ~2.1 Å from a global gold standard FSC of 2.13 Å. Scale bar 2 nm. Density displayed at 5σ.

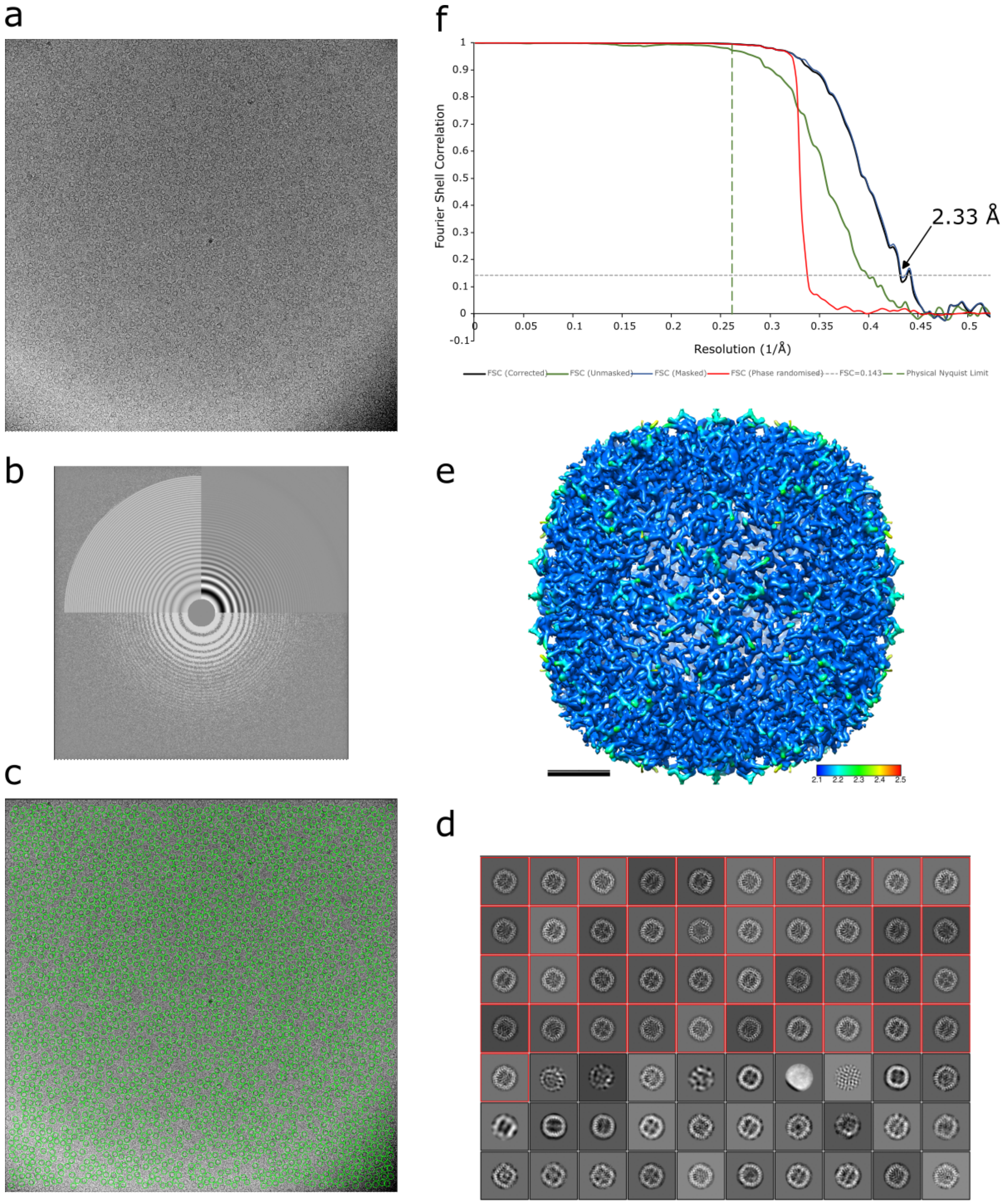


**Figure S3**: Processing of PASR processed, 4K sampled data from second apoferritin dataset. This dataset permits comparison of behaviour with respect to PASR processing for Falcon 4 detectors. (a) Representative micrograph, (b) power spectrum of (a), (c) template picks from (a) are circled in green, (d) result of 2D classification, (e) final reconstruction, coloured by local resolution, extending to a local maximum resolution of ~2.1 Å from a global gold standard FSC of 2.33 Å. Scale bar 2 nm. Density displayed at 5σ.

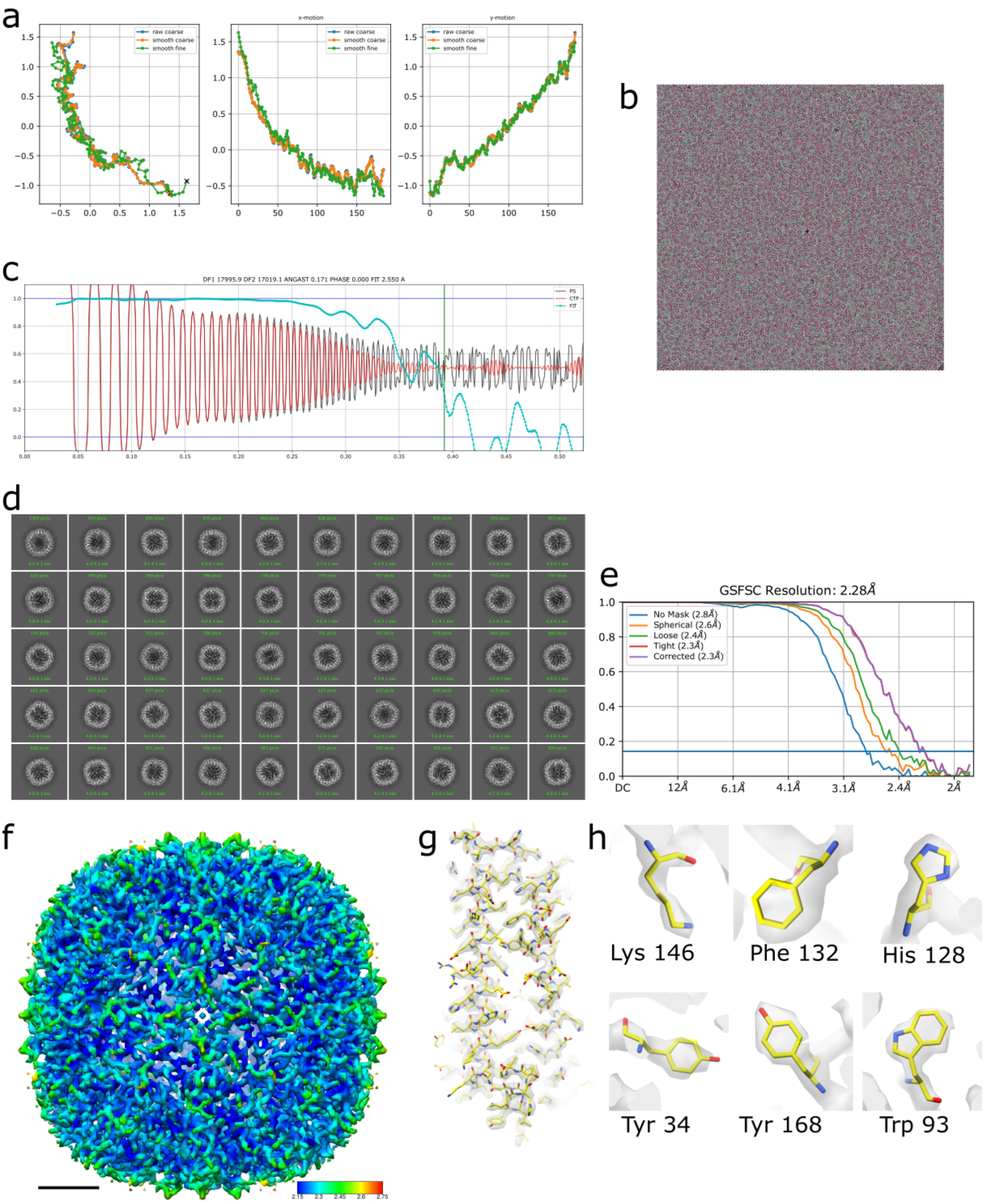


**Figure S4**: CryoSPARC processing of 8K sampling apoferritin dataset. This dataset permits comparison of behaviour with respect to PASR processing for Falcon 4 detectors in CryoSPARC, rather than RELION. (a) Plots of motion during motion correction, (b) representative micrograph with Gaussian blob picking locations in magenta, (c) 1D diagnostic plot of the estimated CTF, (d) representative 2D classes, (e) gold standard FSC output from homogeneous refinement achieving 2.3 Å, (f) 3D reconstruction coloured by local resolution from 2.15-2.75 Å, scale bar equals 2 nm, (g) segmented apoferritin monomer with fitted PDB model (PDBID:7A6A), (h) six representative side chain views. Scale bar equals 2nm.

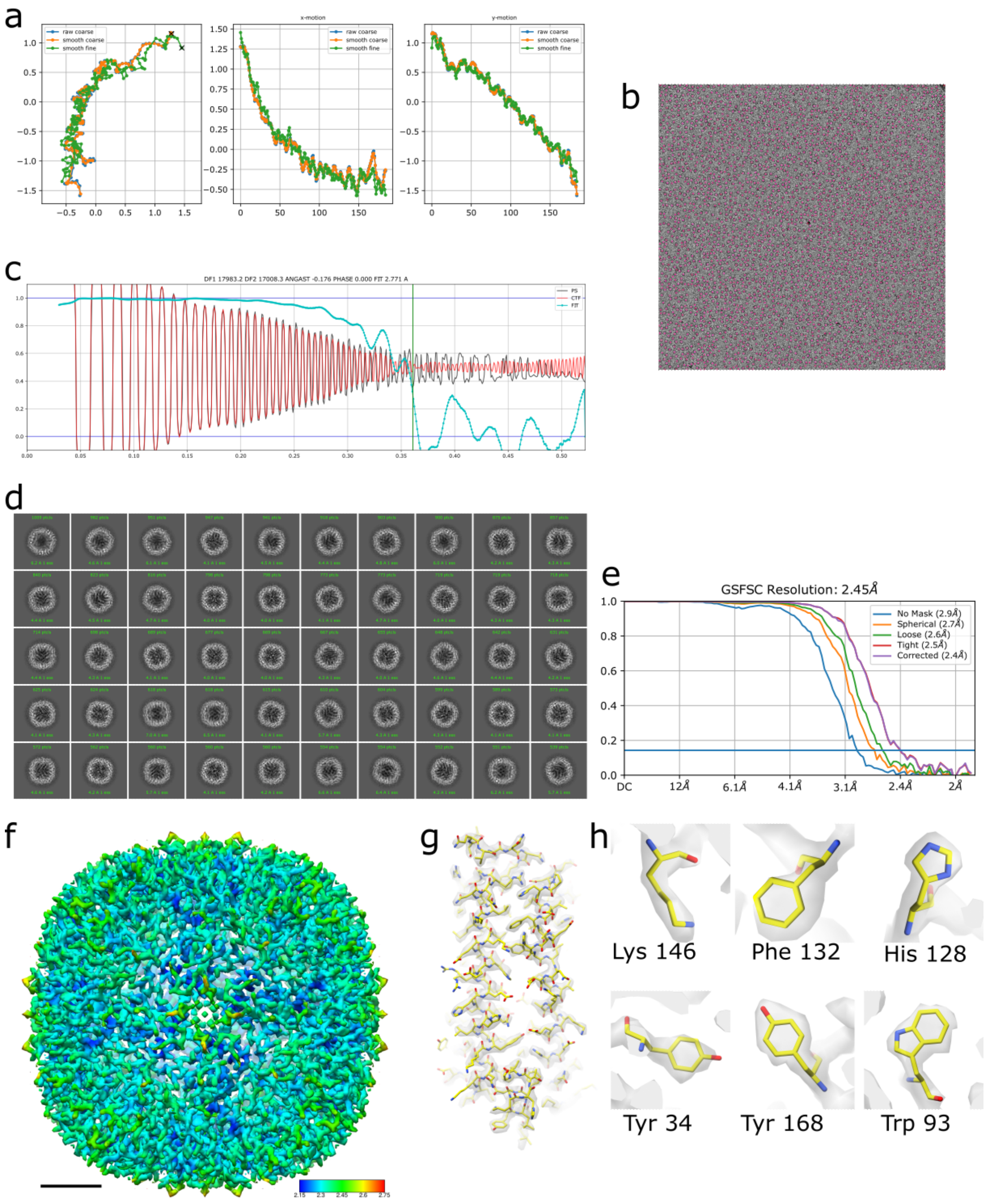


**Figure S5**: CryoSPARC processing of 4K sampling PASR processed apoferritin dataset. This dataset permits comparison of behaviour with respect to PASR processing for Falcon 4 detectors in CryoSPARC, rather than RELION (a) Plots of motion during motion correction. It should be noted that for speed, the PASR micrographs were loaded as MRC format, which reads in with inverted orientation relative to TIFF so the motion plots are flipped in the x-axis, (b) representative micrograph with Gaussian blob picking locations in magenta, (c) 1D diagnostic plot of the estimated CTF, (d) representative 2D classes, (e) gold standard FSC output from homogeneous refinement achieving 2.45 Å, (f) 3D reconstruction coloured by local resolution from 2.15-2.75 Å, scale bar equals 2 nm, (g) segmented apoferritin monomer with fitted PDB model (PDBID:7A6A), (h) six representative side chain views. Scale bar equals 2nm.

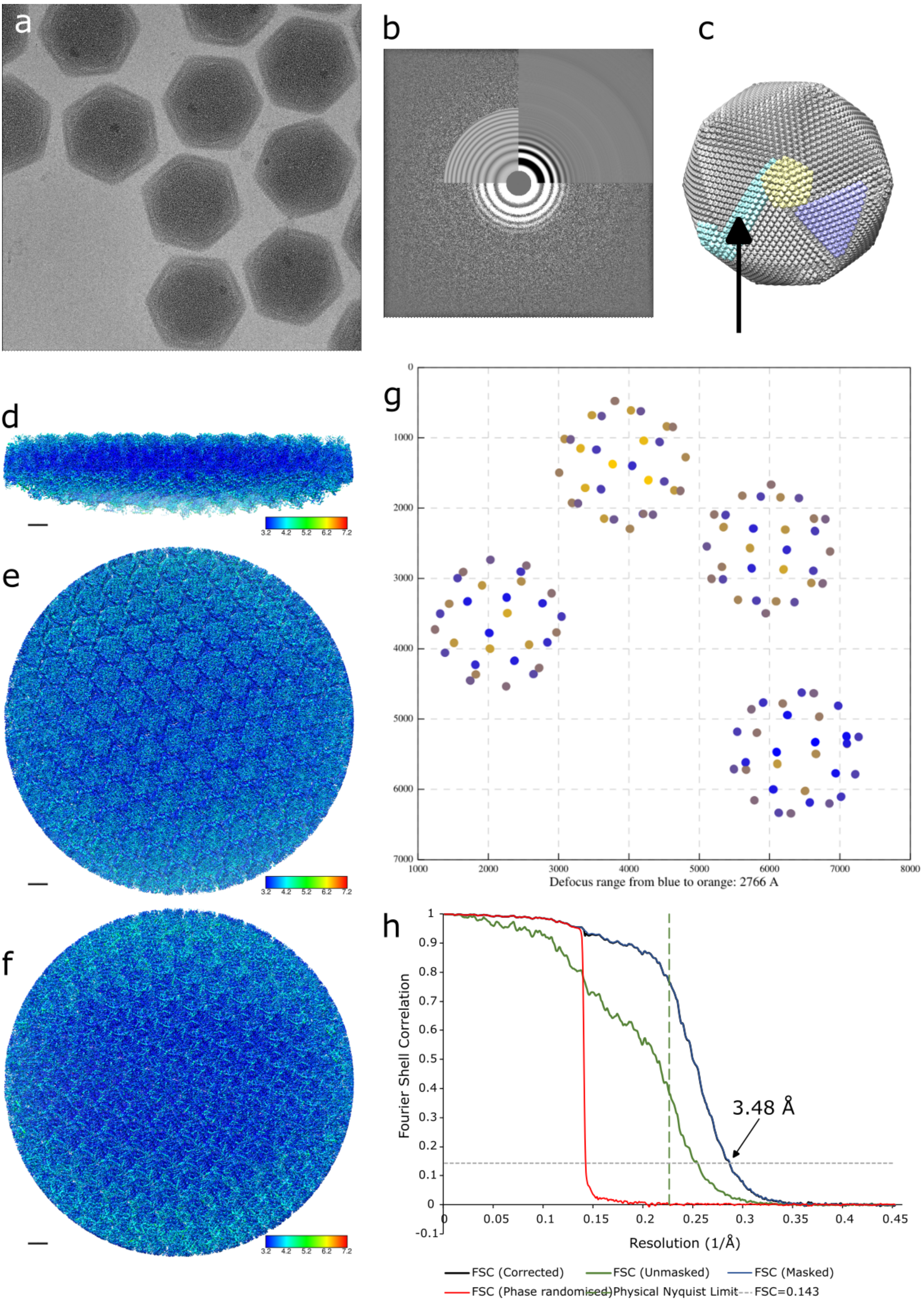
a
b
c
d
e
f
g
h
3.2 4.2 5.2 6.2 7.2
Defocus range from blue to orange: 2766 A
Fourier Shell Correlation
Resolution (1/Å)
3.48 Å
FSC (Corrected)
FSC (Unmasked)
FSC (Masked)
FSC (Phase randomised)
Physical Nyquist Limit
FSC=0.143

**Figure S6**: Twofold block-based reconstruction of PASR processed Melbournevirus. This dataset permits comparison of behaviour with respect to PASR processing for Gatan K2 Summit detectors in counting mode. (a) PASR processed micrograph. (b) Power spectrum and simulated CTF fit from CTFFIND for (a). (c) The whole Melbournevirus with the two, three and fivefold blocks indicated, with an arrow indicating the twofold block. (d) Slice through the centre of the twofold block, showing internal estimated local resolution. (e) view of the outside of the twofold block, coloured by local resolution (f) view of the inside of the twofold block, coloured by local resolution, (g) points in one of the micrographs showing the locations of the twofold block sections, with their local defocus (h) Gold standard Fourier shell correlation of the twofold PASR processed Melbournevirus block, indicating the corrected FSC (black curve), masked FSC (blue curve), unmasked FSC (green curve), phase randomised FSC (red curve), FSC 0.143 metric (dashed grey line) and Nyquist limit from the original data (long dashed green line). Scale bar 5 nm. Density displayed at 3σ.

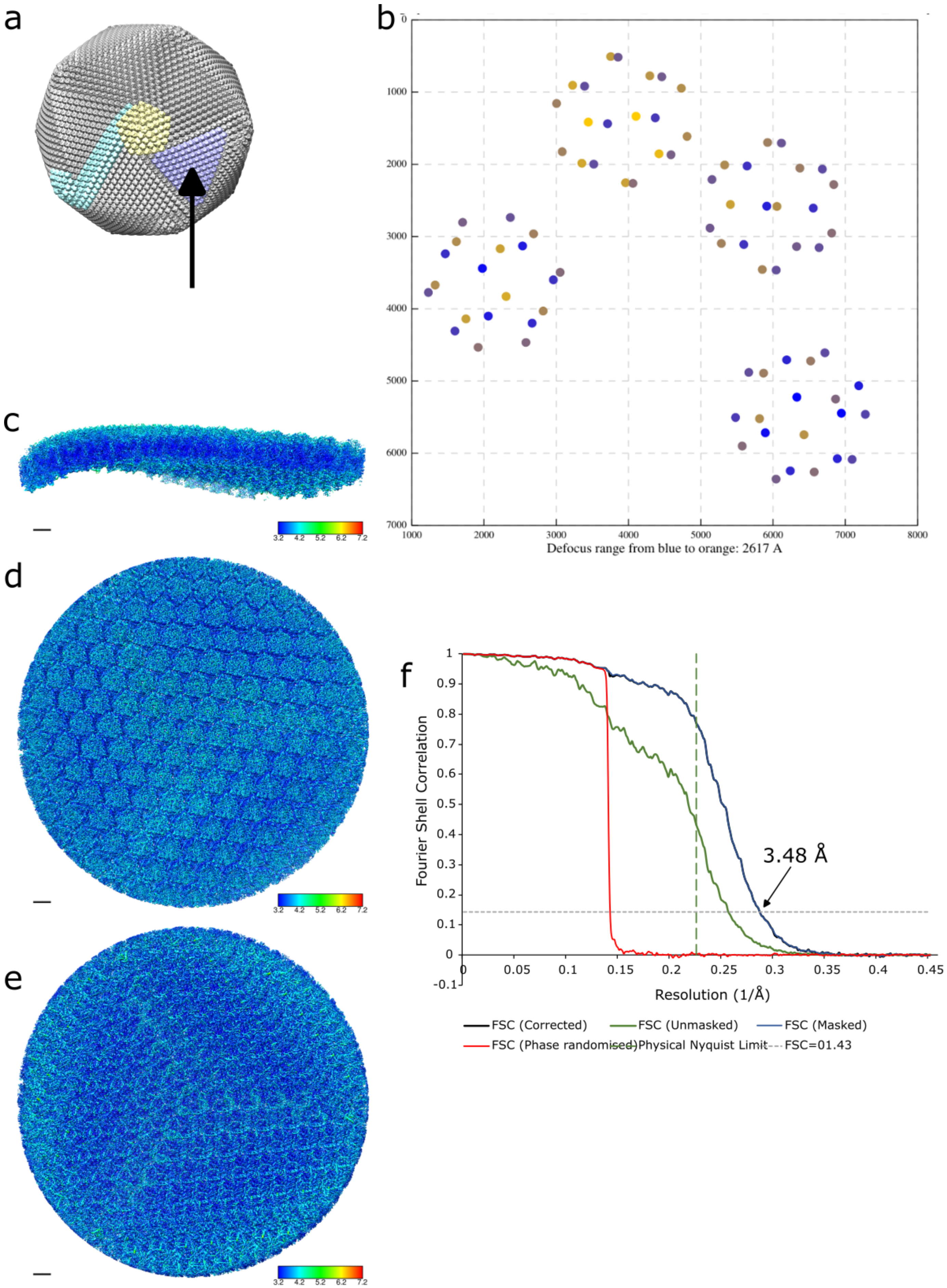
a
b
Defocus range from blue to orange: 2617 A
c
d
e
f
Fourier Shell Correlation
Resolution (1/Å)
3.48 Å
FSC (Corrected)
FSC (Unmasked)
FSC (Masked)
FSC (Phase randomised)
Physical Nyquist Limit
FSC=01.43

**Figure S7**: Threefold block-based reconstruction of PASR processed Melbournevirus. This dataset permits comparison of behaviour with respect to PASR processing for Gatan K2 Summit detectors in counting mode. (a) The whole Melbournevirus with the two, three and fivefold blocks indicated, with an arrow indicating the threefold block. (b) points in one of the micrographs showing the locations of the threefold block sections, with their local defocus, (c) Slice through the centre of the threefold block, showing internal estimated local resolution. (d) view of the outside of the threefold block, coloured by local resolution (e) view of the inside of the threefold block, coloured by local resolution, (f) Gold standard Fourier shell correlation of the threefold PASR processed Melbournevirus block, indicating the corrected FSC (black curve), masked FSC (blue curve), unmasked FSC (green curve), phase randomised FSC (red curve), FSC 0.143 metric (dashed grey line) and Nyquist limit from the original data (long dashed green line). Scale bar 5 nm. Density displayed at 3σ.

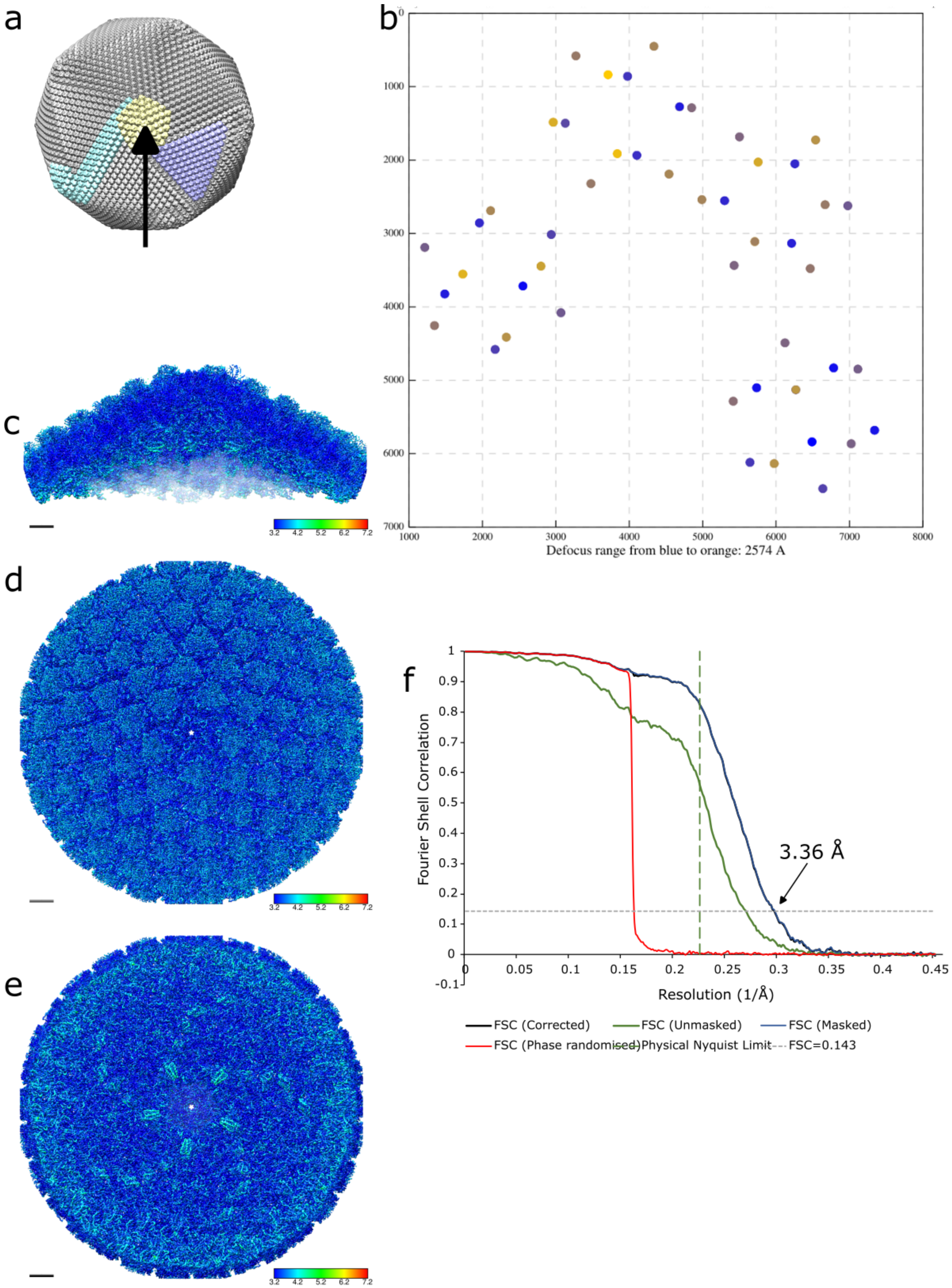

a
b
c
d
e
f
1000 2000 3000 4000 5000 6000 7000 8000
0 1000 2000 3000 4000 5000 6000 7000
Defocus range from blue to orange: 2574 A
3.2 4.2 5.2 6.2 7.2
Fourier Shell Correlation
Resolution (1/Å)
3.36 Å
FSC (Corrected)
FSC (Unmasked)
FSC (Masked)
FSC (Phase randomised)
Physical Nyquist Limit
FSC=0.143

**Figure S8**: Fivefold block-based reconstruction of PASR processed Melbournevirus. This dataset permits comparison of behaviour with respect to PASR processing for Gatan K2 Summit detectors in counting mode. (a) The whole Melbournevirus with the two, three and fivefold blocks indicated, with an arrow indicating the fivefold block. (b) points in one of the micrographs showing the locations of the fivefold block sections, with their local defocus, (c) Slice through the centre of the fivefold block, showing internal estimated local resolution. (d) view of the outside of the fivefold block, coloured by local resolution (e) view of the inside of the fivefold block, coloured by local resolution, (f) Gold standard Fourier shell correlation of the threefold PASR processed Melbournevirus block, indicating the corrected FSC (black curve), masked FSC (blue curve), unmasked FSC (green curve), phase randomised FSC (red curve), FSC 0.143 metric (dashed grey line) and Nyquist limit from the original data (long dashed green line). Scale bar 5 nm. Density displayed at 3σ.

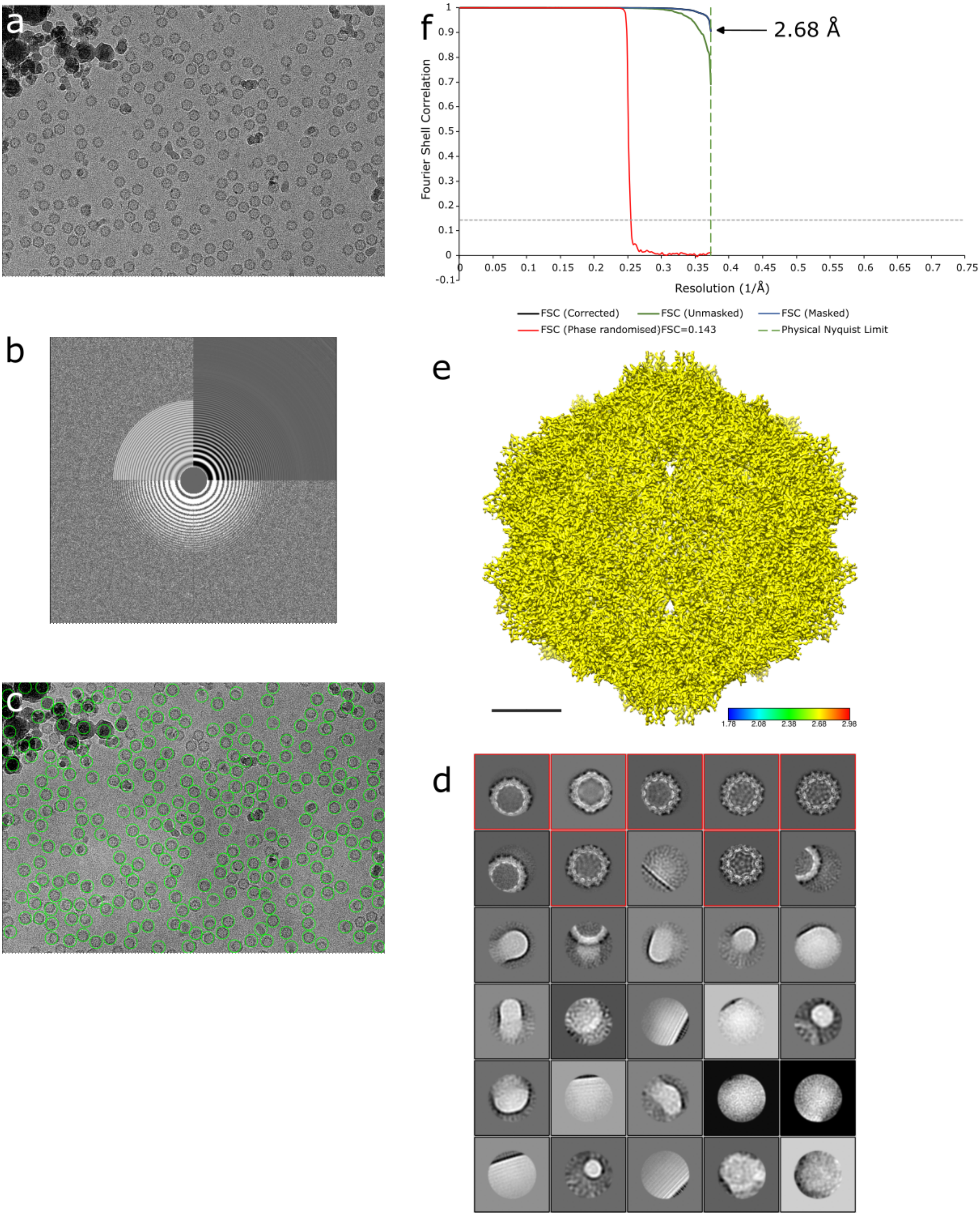


**Figure S9**: Processing of the original adeno-associated virus dataset. This dataset permits comparison of behaviour with respect to PASR processing for Gatan K3 detectors in correlative double sampling (CDS) mode. (a) Representative micrograph, (b) power spectrum showing good fit to Thon rings for (a), (c) LoG autopicking of (a), (d) representative 2D classes, with selected classes highlighted in red, (e) final reconstruction, coloured by local resolution at 2.68 Å, scale bar equals 5 nm, (f) RELION gold standard FSC, showing the original data reaching the Nyquist limit.

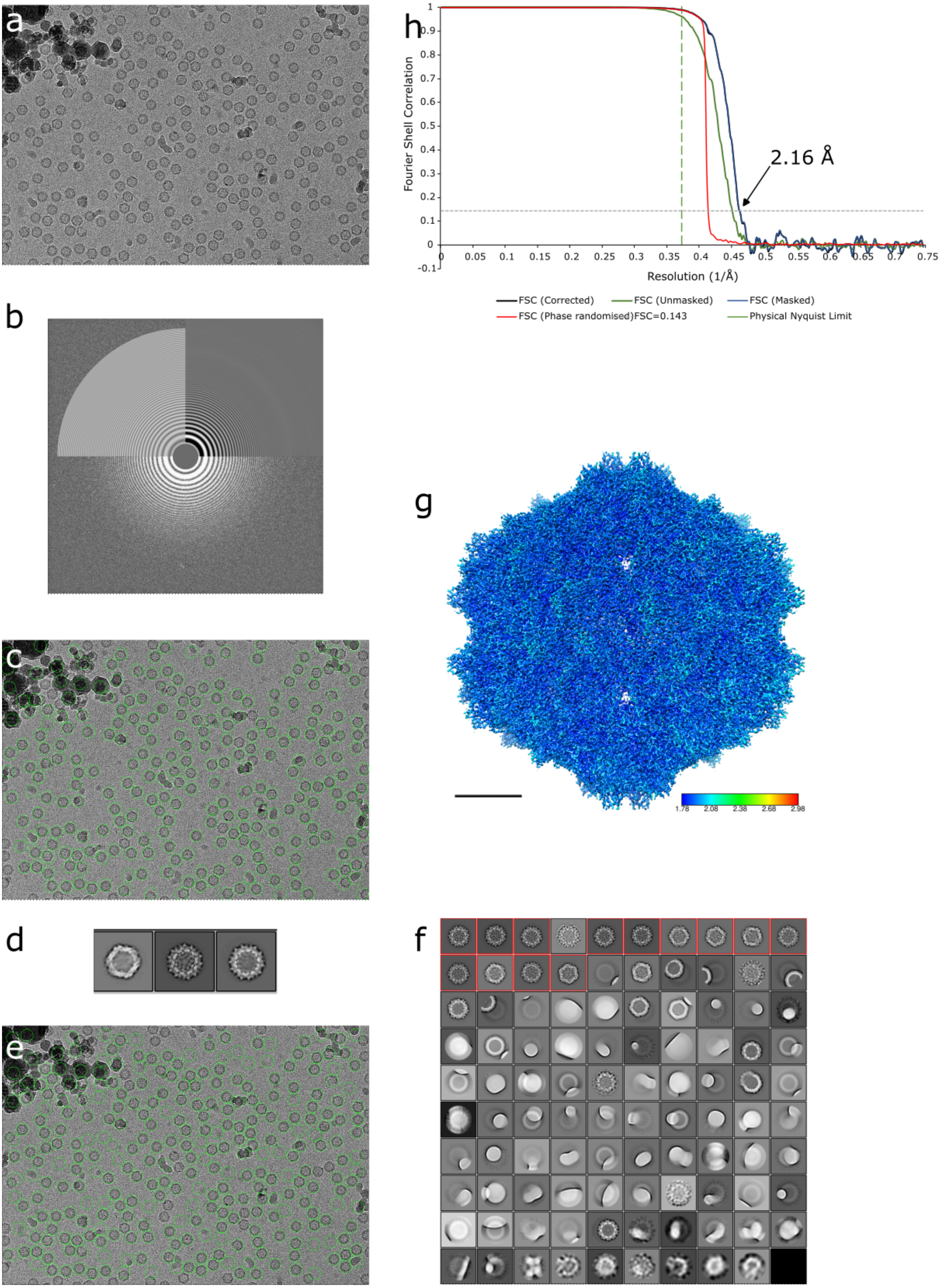
a
b
c
d
e
f
g
h
Fourier Shell Correlation
Resolution (1/Å)
2.16 Å
1
0.9
0.8
0.7
0.6
0.5
0.4
0.3
0.2
0.1
0
-0.1
0.05
0.1
0.15
0.2
0.25
0.3
0.35
0.4
0.45
0.5
0.55
0.6
0.65
0.7
0.75
FSC (Corrected)
FSC (Unmasked)
FSC (Masked)
FSC (Phase randomised)FSC=0.143
Physical Nyquist Limit
1.78
2.08
2.38
2.68
2.98

**Figure S10**: Processing of the PASR processed adeno-associated virus dataset. This dataset permits comparison of behaviour with respect to PASR processing for Gatan K3 detectors in correlative double sampling (CDS) mode. (a) Representative micrograph, (b) power spectrum showing good fit to Thon rings for (a), (c) LoG autopicking of (a), (d) representative 2D classes, which were used for template autopicking, (e) micrograph with template autopicks circled in green, (f) 2D classes, with selected classes highlighted in red, (g) final reconstruction, coloured by local resolution, extending to ~1.78 Å, scale bar equals 5 nm, (h) RELION gold standard FSC, showing the PASR processed data exceeding the original Nyquist limit.

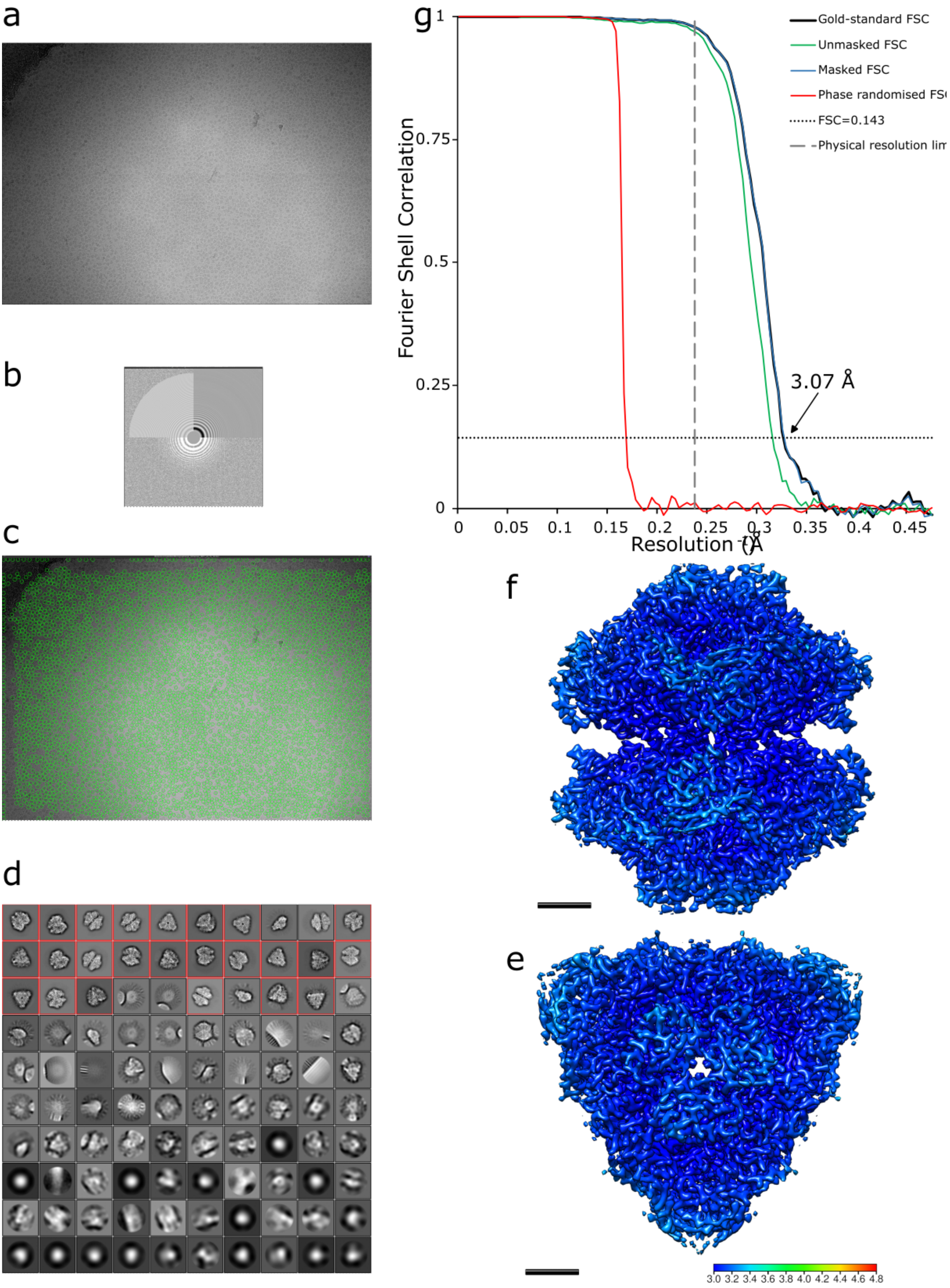

a
b
c
d
g
Gold-standard FSC
Unmasked FSC
Masked FSC
Phase randomised FS
FSC=0.143
Physical resolution li
Fourier Shell Correlation
1
0.75
0.5
0.25
0
3.07 Å
0
0.05
0.1
0.15
0.2
0.25
0.3
0.35
0.4
0.45
Resolution (Å⁻¹)
f
e
3.0 3.2 3.4 3.6 3.8 4.0 4.2 4.4 4.6 4.8

**Figure S11**: Processing of jack bean urease (EMPIAR-10549). This dataset permits comparison of behaviour with respect to PASR processing for Gatan K3 detectors in super resolution mode, and for 200 kV acceleration voltage microscopes. (a) a representative micrograph, (b) power spectrum of (a), (c) LoG autopicking of (a), note that the LoG autopicking algorithm "mis-picks" at the edges, (d) 2D classes, with selected classes highlighted in red, (e) side view of jack bean urease reconstruction, coloured by local resolution, with resolution extending to ~3 Å, (f) side view of (e). (g) RELION gold standard FSC showing the data reaching the physical Nyquist limit. Scale bar equals 2 nm.

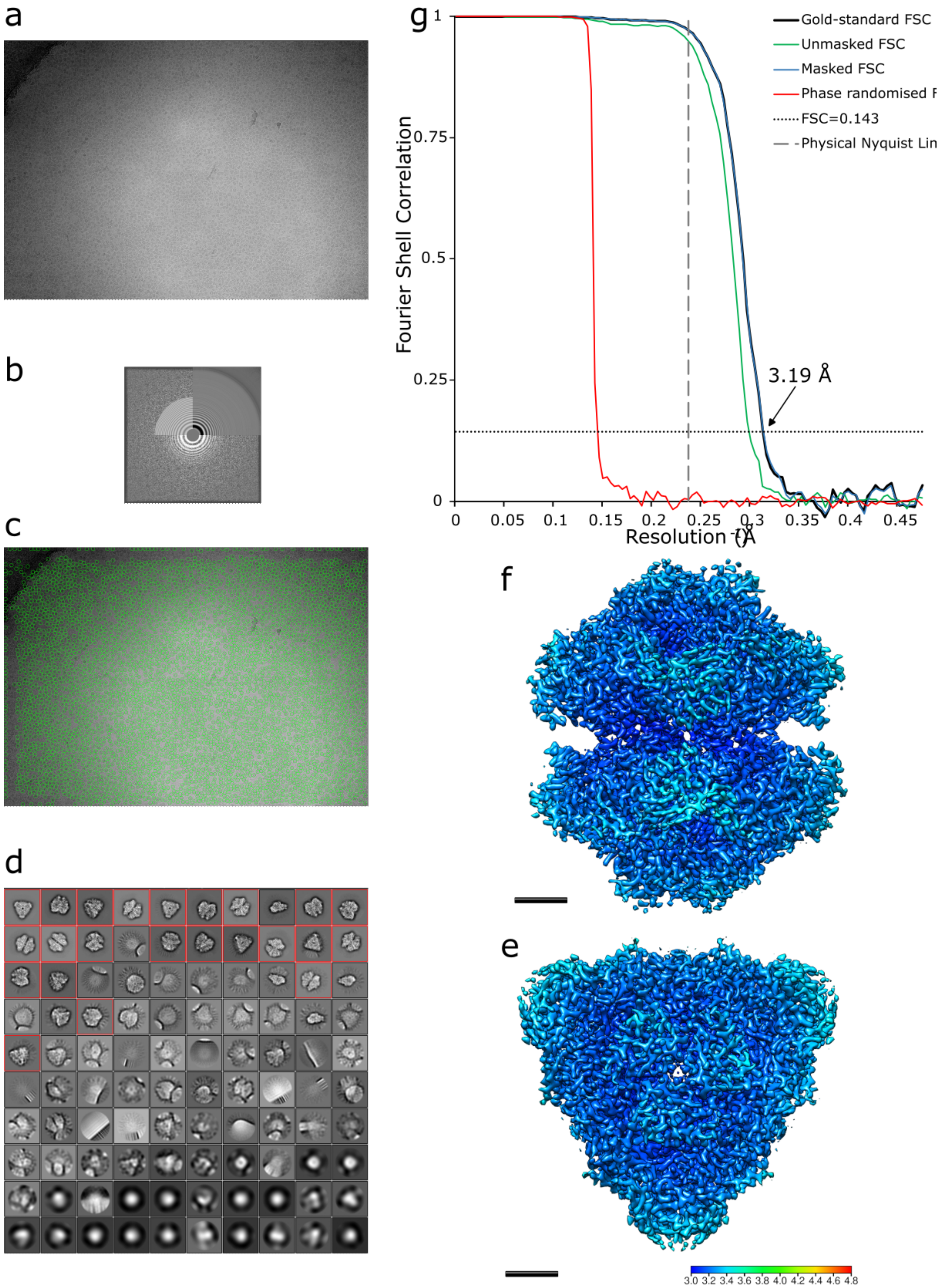

a
b
c
d
g
1
0.75
0.5
0.25
0
Fourier Shell Correlation
0
0.05
0.1
0.15
0.2
0.25
0.3
0.35
0.4
0.45
Resolution (Å)
Gold-standard FSC
Unmasked FSC
Masked FSC
Phase randomised F
FSC=0.143
Physical Nyquist Lin
3.19 Å
f
e
3.0 3.2 3.4 3.6 3.8 4.0 4.2 4.4 4.6 4.8

**Figure S12**: Processing of jack bean urease (EMPIAR-10549) after PASR processing. This dataset permits comparison of behaviour with respect to PASR processing for Gatan K3 detectors in super resolution mode, and for 200 kV acceleration voltage microscopes. (a) a representative micrograph, (b) power spectrum of (a), (c) LoG autopicking of (a), note that the LoG autopicking algorithm "mis-picks" at the edges, (d) 2D classes, with selected classes highlighted in red, (e) side view of jack bean urease reconstruction, coloured by local resolution, with resolution extending to ~3 Å, (f) side view of (e). (g) RELION gold standard FSC showing the data reaching the physical Nyquist limit. Scale bar equals 2 nm.

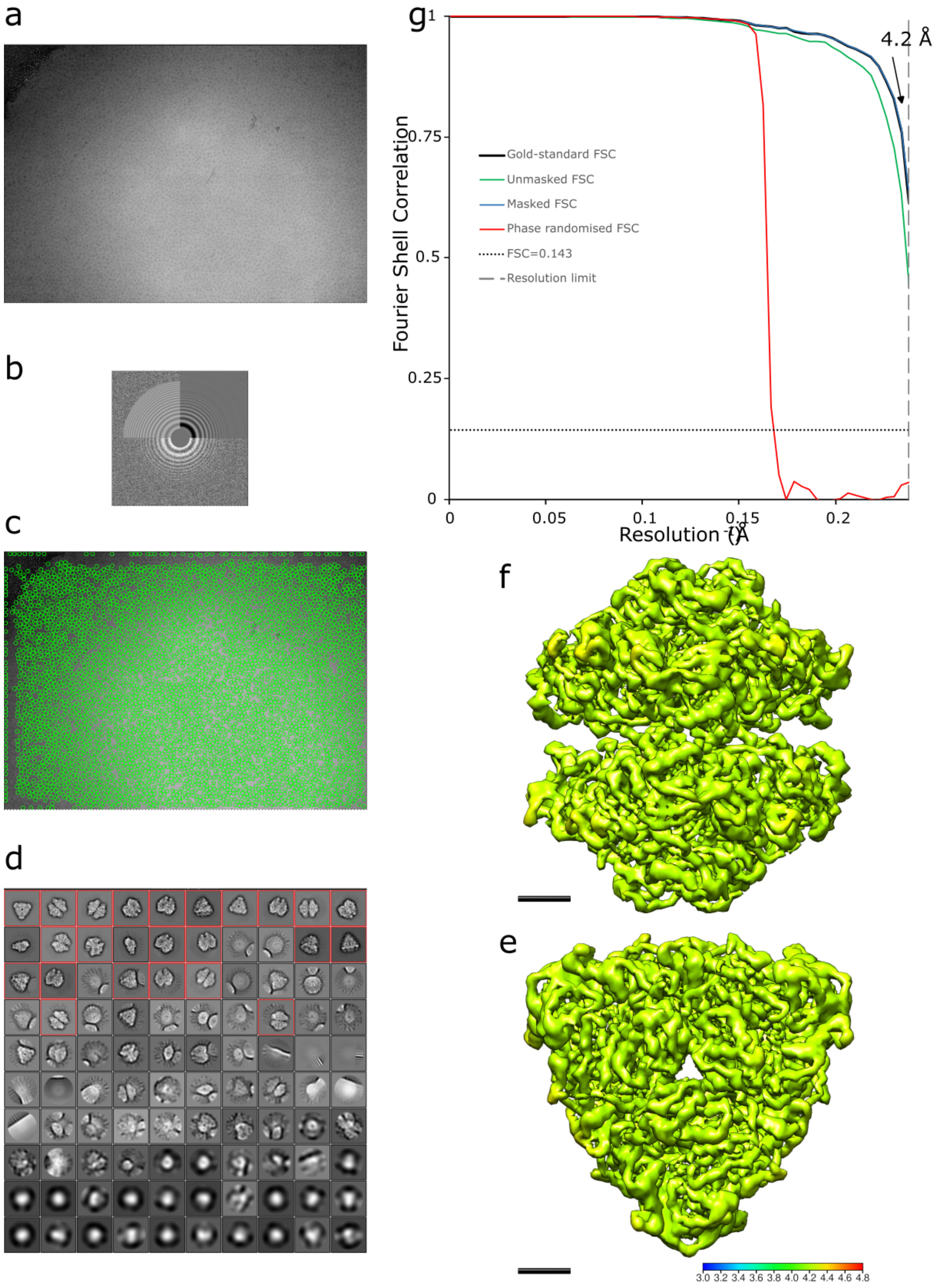
a
b
c
d
g
Fourier Shell Correlation
Resolution (Å⁻¹)
4.2 Å
Gold-standard FSC
Unmasked FSC
Masked FSC
Phase randomised FSC
FSC=0.143
Resolution limit
0
0.25
0.5
0.75
1
0.05
0.1
0.15
0.2
f
e
3.0 3.2 3.4 3.6 3.8 4.0 4.2 4.4 4.6 4.8

**Figure S13**: Processing of jack bean urease (EMPIAR-10549) after binning micrographs to physical Nyquist. This dataset permits comparison of behaviour with respect to PASR processing for Gatan K3 detectors in super resolution mode, and for 200 kV acceleration voltage microscopes. (a) a representative micrograph, (b) power spectrum of (a), (c) LoG autopicking of (a), note that the LoG autopicking algorithm "mis-picks" at the edges, (d) 2D classes, with selected classes highlighted in red, (e) side view of jack bean urease reconstruction, coloured by local resolution (structure is globally 4.2 Å), (f) side view of (e). (g) RELION gold standard FSC showing the data reaching the physical Nyquist limit. Scale bar equals 2 nm.

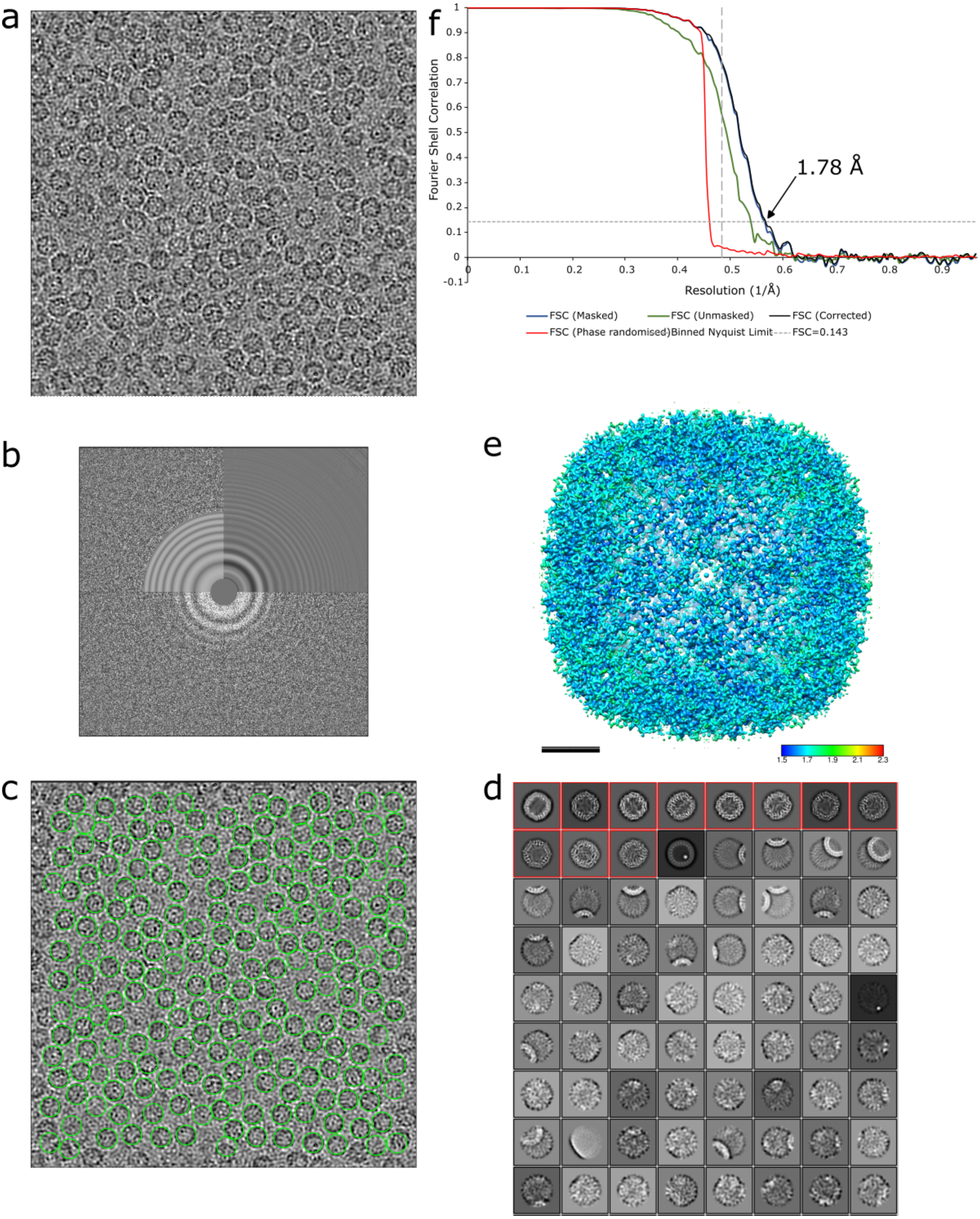


**Figure S14**: Application of PASR pre-motion correction (i.e., to movies) on EMPIAR-10216. This dataset permits comparison of behaviour with respect to PASR processing for Falcon 3 detectors, which do not have a super resolution mode. (a) Representative micrograph, (b) power spectrum of (a), (c) template picks from (a) are circled in green, (d) result of 2D classification, (e) final reconstruction, coloured by local resolution, extending to a local maximum resolution of ~1.5 Å from a global gold standard FSC of 1.78 Å. Scale bar 2 nm. Density displayed at 5σ.

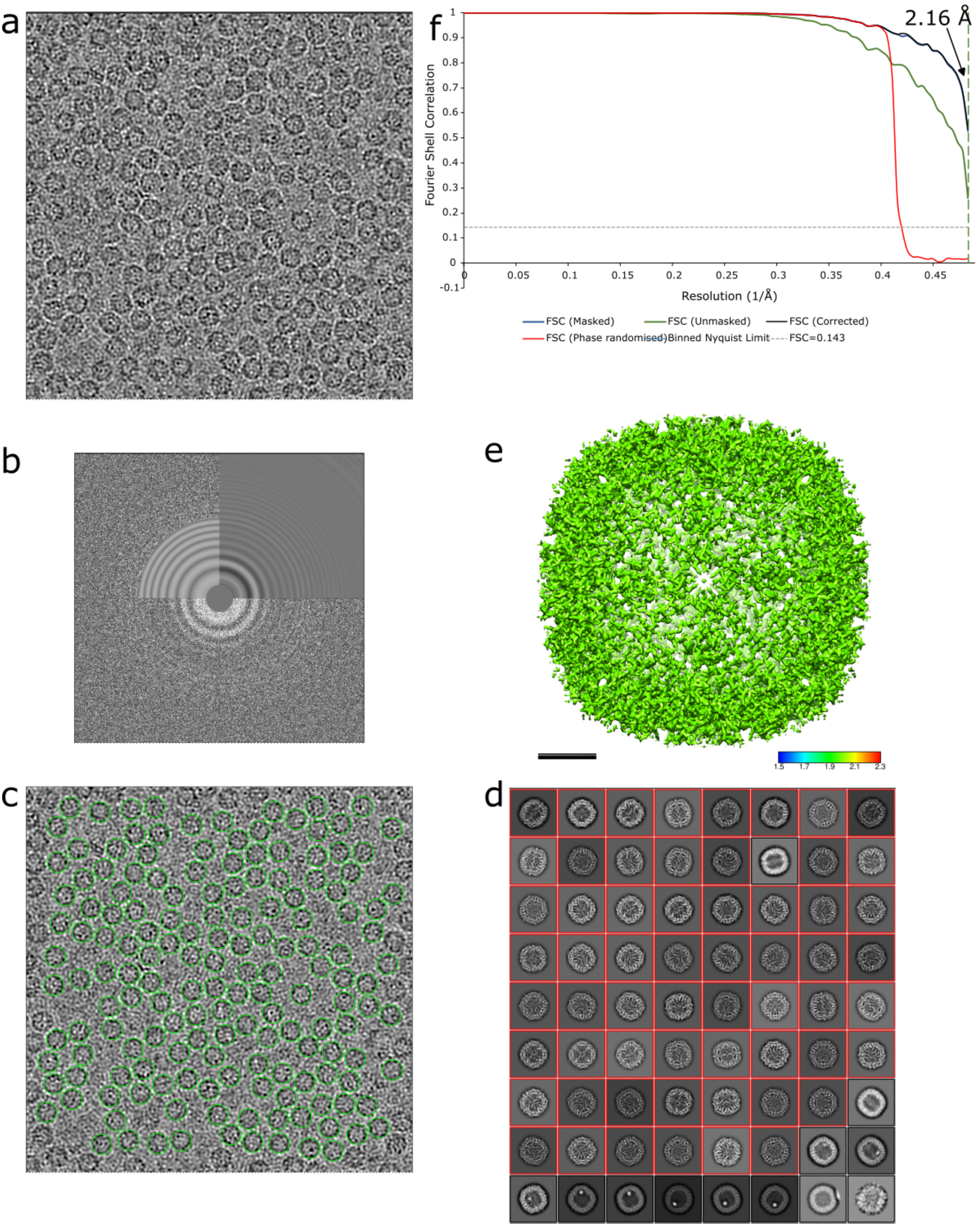


**Figure S15**: Binned processing of EMPIAR-10216. This dataset permits comparison of behaviour with respect to PASR processing for Falcon 3 detectors, which do not have a super resolution mode. (a) Representative micrograph, (b) power spectrum of (a), (c) template picks from (a) are circled in green, (d) result of 2D classification, (e) final reconstruction, coloured by local resolution, with the entire reconstruction estimated at the binned Nyquist limit (2.068 Å) from a global gold standard FSC of 2.068 Å. Scale bar 2 nm. Density displayed at 5σ.

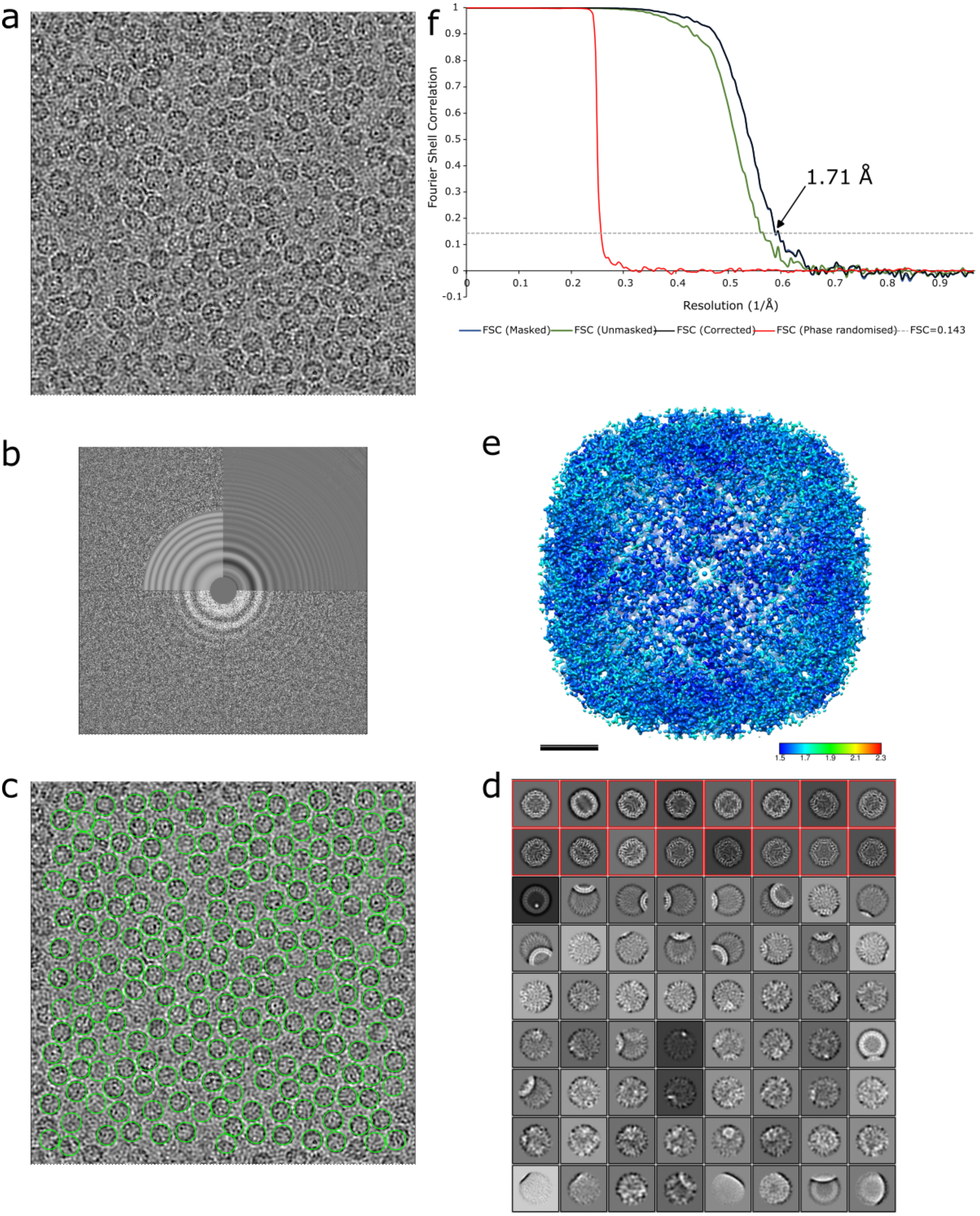


**Figure S16**: Reprocessing of the original EMPIAR-10216 dataset. This dataset permits comparison of behaviour with respect to PASR processing for Falcon 3 detectors, which do not have a super resolution mode. (a) Representative micrograph, (b) power spectrum of (a), (c) template picks from (a) are circled in green, (d) result of 2D classification, (e) final reconstruction, coloured by local resolution, extending to a local maximum resolution of ~1.5 Å from a global gold standard FSC of 1.71 Å. Scale bar 2 nm. Density displayed at 5σ.

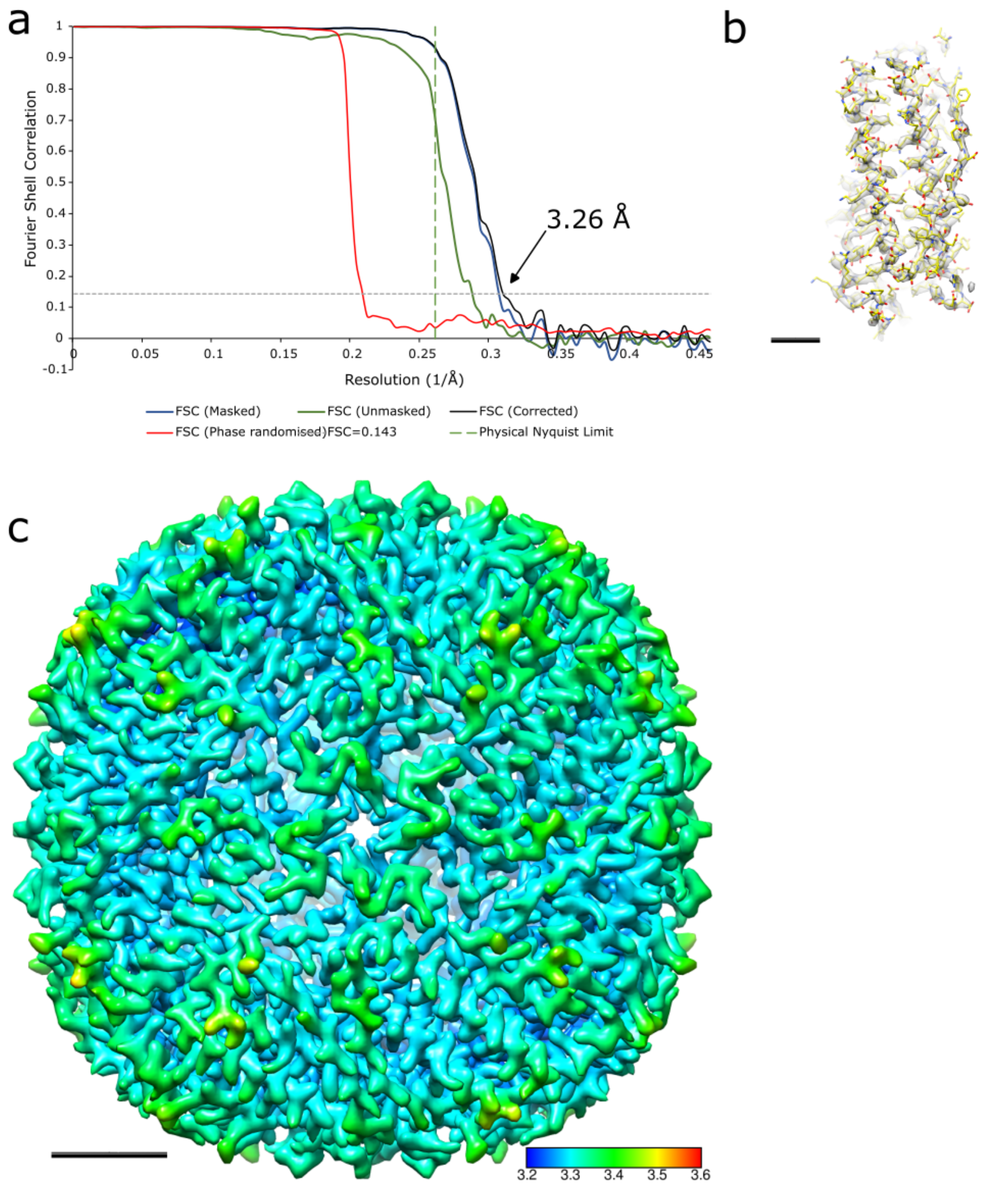


**Figure S17**: Application of PASR post-motion correction (i.e., to single frame micrographs) on the second apoferritin (64,000×) dataset. This dataset permits comparison of behaviour with respect to PASR processing for single frame micrograph data for smaller particles. (a) RELION gold standard FSC. The corrected curve (black line) falls to 3.26 Å at FSC = 0.143. (b) apoferritin monomer, with fitted PDB showing what would be expected at ~3.3 Å; side chain density but lacking clear features. (c) post-motion correction PASR reconstruction with resolution only extending to ~3.2 Å, rather than the ~2.1 Å if applying PASR prior to motion correction. Scale bar for (b) equals 1 nm, for (c) equals 2 nm.

a

b

c

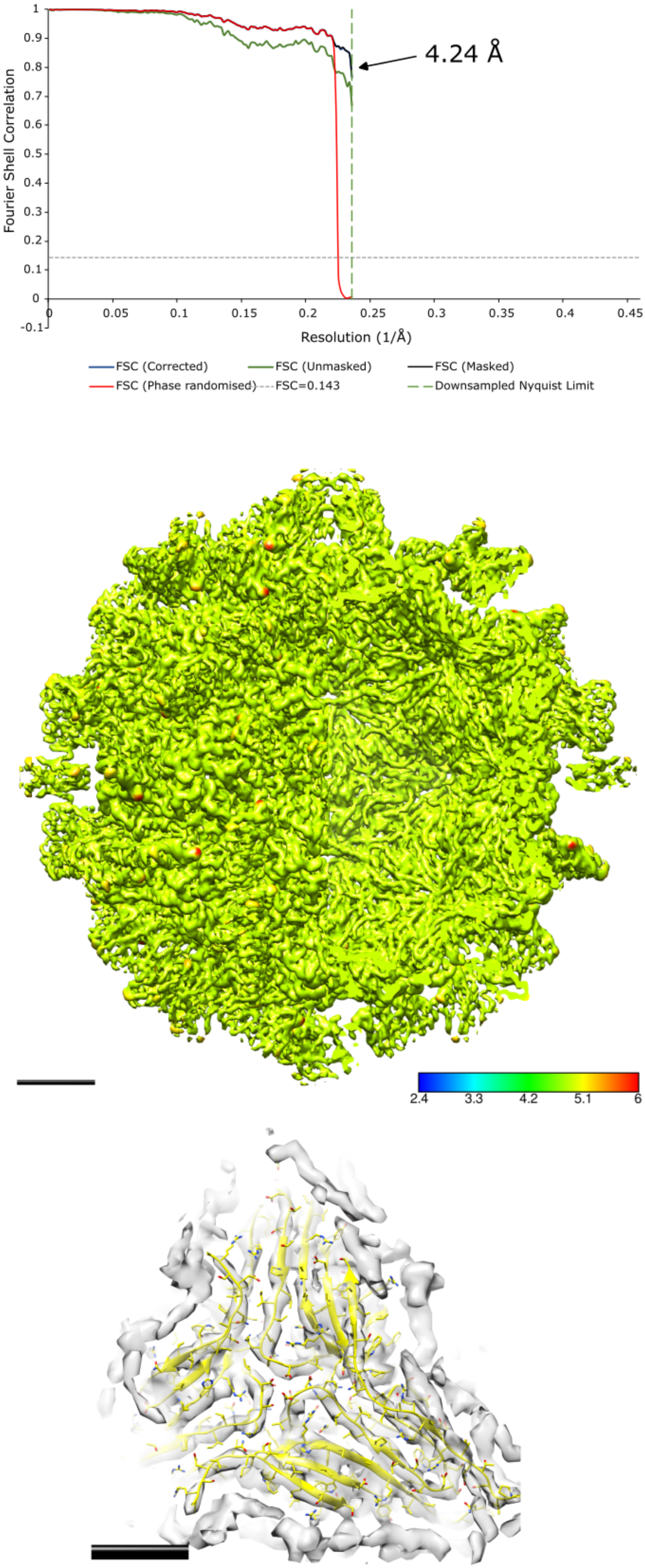

4.24 Å
Fourier Shell Correlation
Resolution (1/Å)
1
0.9
0.8
0.7
0.6
0.5
0.4
0.3
0.2
0.1
0
-0.1
0
0.05
0.1
0.15
0.2
0.25
0.3
0.35
0.4
0.45
FSC (Corrected)
FSC (Unmasked)
FSC (Masked)
FSC (Phase randomised)
FSC=0.143
Downsampled Nyquist Limit
2.4
3.3
4.2
5.1
6

**Figure S18**: Nodavirus (EMPIAR-10203) downsampled before PASR processing. This dataset permits comparison of behaviour with respect to PASR processing for single frame micrographs which have already undergone motion correction, however it is not recommended. (a) RELION gold standard FSC curve demonstrating the downsampled Nyquist limit, (b) binned reconstruction, coloured by local resolution showing almost the entire map is at the Nyquist limit, scale bar 5 nm. Density displayed at 3σ. (c) Focussed view of the internal surface of the capsid, scale bar 2 nm. Density displayed at 6σ.

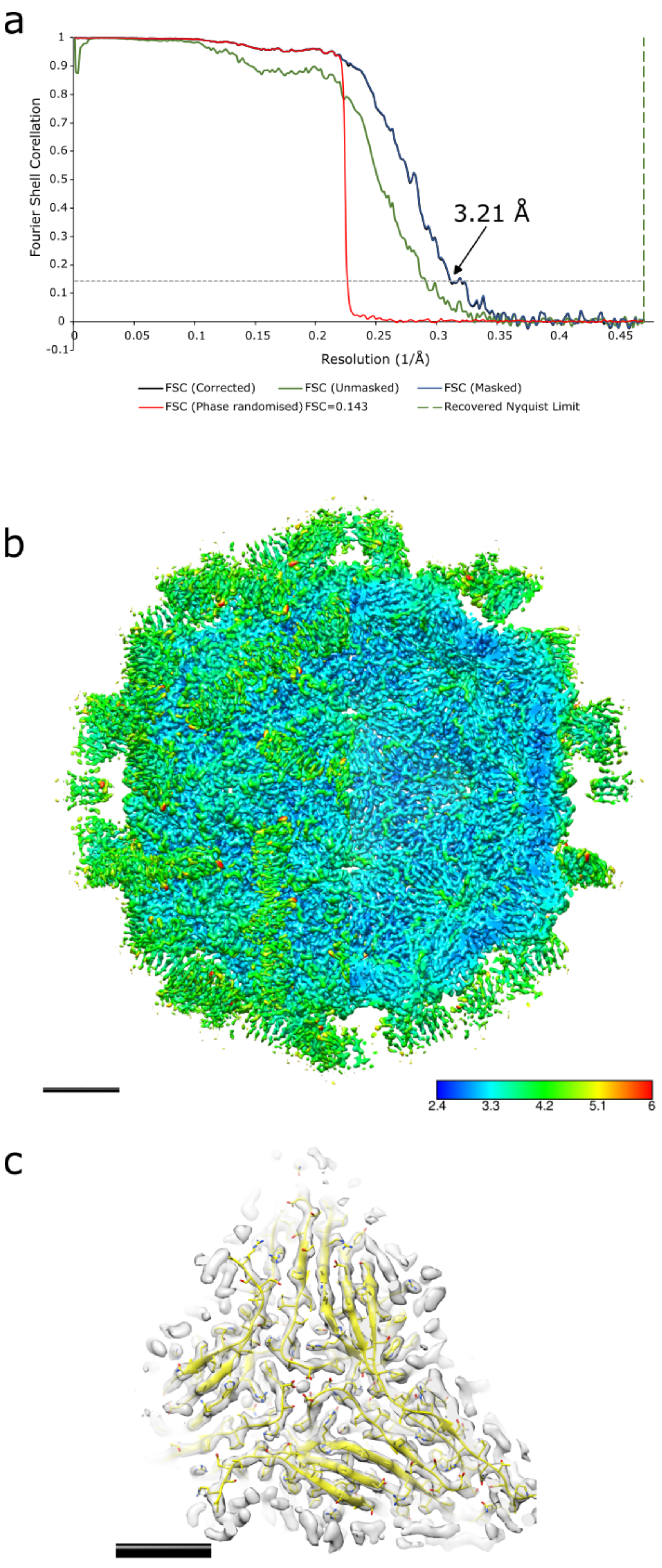
a
Fourier Shell Corellation
Resolution (1/Å)
3.21 Å
FSC (Corrected)
FSC (Unmasked)
FSC (Masked)
FSC (Phase randomised)
FSC=0.143
Recovered Nyquist Limit
b
2.4
3.3
4.2
5.1
6
c

**Figure S19**: Nodavirus (EMPIAR-10203) after downsampled then upscaled with PASR processing. This dataset permits comparison of behaviour with respect to PASR processing for single frame micrographs which have already undergone motion correction, however it is not recommended. (a) RELION gold standard FSC curve, reaching 3.21 Å, (b) PASR reconstruction, coloured by local resolution which extends to ~2.4 Å, scale bar 5 nm. Density displayed at 3σ. (c) Focussed view of the internal surface of the capsid, scale bar 2 nm. Density displayed at 6σ.

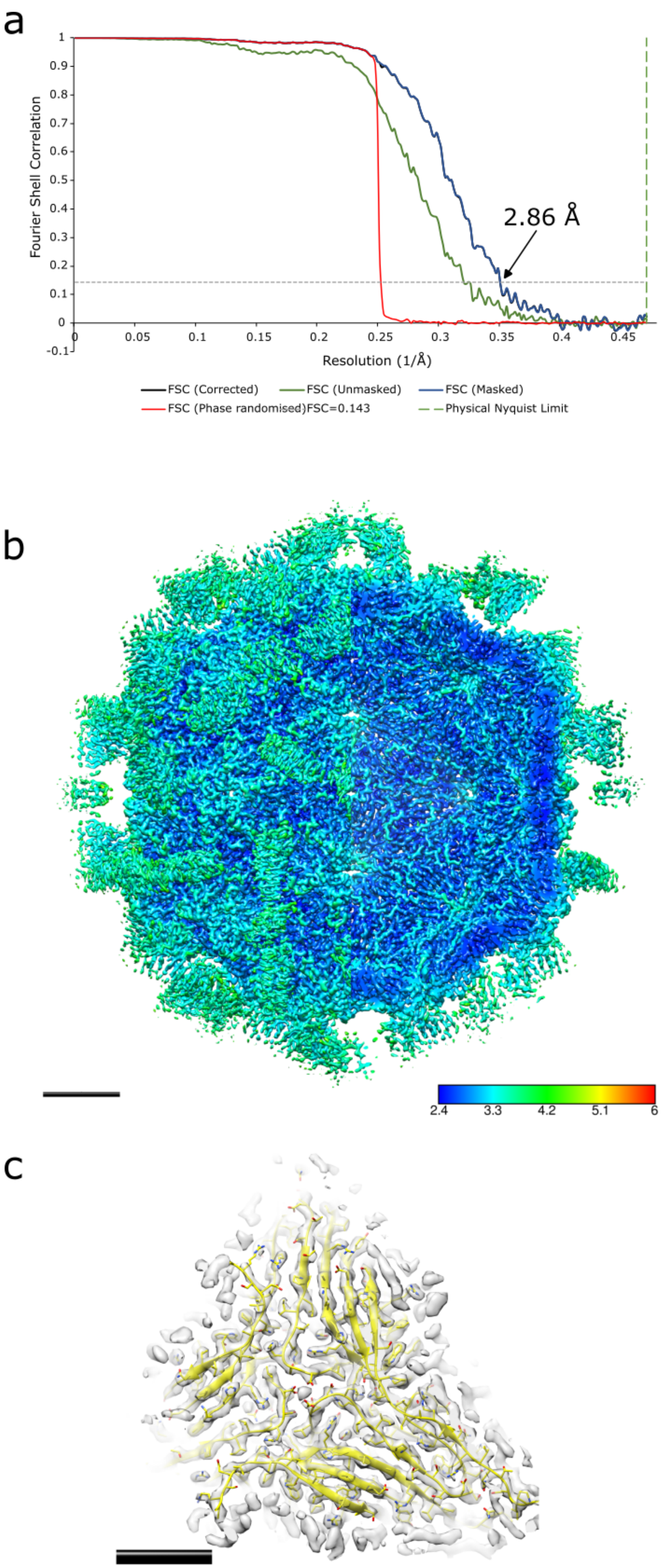

a
Fourier Shell Correlation
Resolution (1/Å)
2.86 Å
FSC (Corrected)
FSC (Unmasked)
FSC (Masked)
FSC (Phase randomised)FSC=0.143
Physical Nyquist Limit
b
2.4
3.3
4.2
5.1
6
c

**Figure S20**: Nodavirus (EMPIAR-10203) original data reprocessed for comparison with a modern cryo-EM processing suite. This dataset permits comparison of behaviour with respect to PASR processing for single frame micrographs which have already undergone motion correction, however it is not recommended. (a) RELION gold standard FSC curve, reaching 2.86 Å, (b) Reconstruction of original data, coloured by local resolution with extends to ~2.4 Å, scale bar 5 nm. Density displayed at 3σ. (c) Scale bar 2 nm. Density displayed at 6σ.

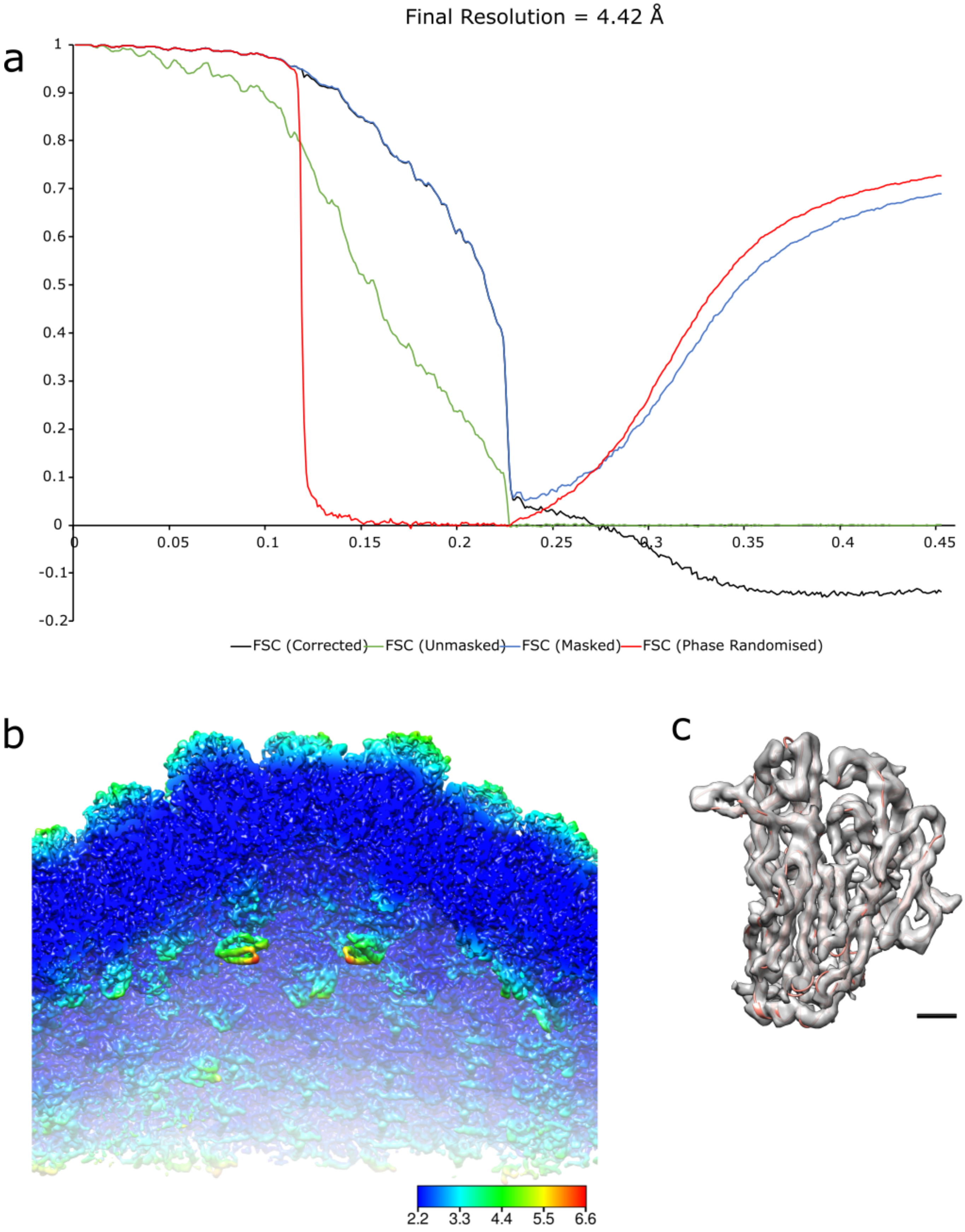


**Figure S21**: The effect of zero padding on the half maps of original data, demonstrating that PASR is acting as if acquired with super resolution. (a) RELION gold standard FSC, showing immediately that zero padding is harmful; while the unmasked half map FSC acts as expected, the mask, corrected and phase randomised FSCs display behaviour which should immediately alert the user that something is awry, (b) sliced view, coloured by estimated local resolution from the zero padded half maps, estimating resolution extending to the zero padded sampling limit. Map features do not support this resolution estimate and appear to be the same as the original map of 4.4 Å. (c) extracted major capsid protein monomer showing complete lack of features one would expect from a 2.2 Å map. Scale bar equals 1 nm.
